# Projection populations form a dorsoventral gradient of reward representations in the lateral septum

**DOI:** 10.64898/2026.10.20.760513

**Authors:** Jennifer Isaac, Sonia Karkare, Hymavathy Balasubramanian, Viktor Olah, Matthew Rowan, Malavika Murugan

## Abstract

Several decades ago, the septal region was identified as the most dependable location of intracranial self-stimulation (ICSS) in the brain, but the neural substrates that mediate septal self-stimulation are still unknown. One part of the septal region, the lateral septum (LS), contributes to a number of motivated behaviors and may promote reward-seeking. However, the LS contains different anatomical subregions, complicating the characterization of its reward representations. We examine the responses of LS neurons along its dorsoventral axis using cellular resolution calcium imaging during an operant assay. We find that a greater proportion of neurons are excited in response to sucrose reward in the dorsal LS, while a greater proportion are inhibited in response to sucrose reward in the ventral LS. Additionally, we identify two distinct LS projection populations, one to the nucleus accumbens (NAc) and another to the bed nucleus of the stria terminalis (BNST), which are located in the dorsal and ventral regions of the LS, respectively. These two projection populations also have opposing reward responses and their corresponding optogenetic manipulation supports ICSS. These findings demonstrate how LS projection populations contribute to its functional heterogeneity, especially as it pertains to reward-seeking behavior.

## Introduction

Since it was initially shown that the septal region produces the most reliably reinforcing self-stimulation compared to other brain regions, researchers have sought to determine the exact source of positive reinforcement within the septal region^1^. These early studies used electrodes that spanned the entirety of the septal region which consists of the lateral septum (LS), medial septum, diagonal band of broca and other fibers of passage, making it difficult to ascertain the location of self-stimulation^2^. Additional studies that reconstructed the location of self-stimulation electrodes in the LS found that self-stimulation was elicited at all positions along the anatomical axes of the LS, independent of proximity to dopamine terminals in the LS^3^. This led researchers to term the LS the “elusive substrate” of self-stimulation^3^.

Around the time that the septal region was established as a center of rewarding self-stimulation, it was also found that lesions to the septal region caused a hyper-aggressive phenotype known as septal rage^4–6^. Interestingly, septal lesions that resulted in septal rage also abolished self-stimulation of other canonical reward centers in the brain such as the ventral tegmental area (VTA)^7^. This finding suggests that the LS is broadly involved in reward-related behaviors beyond its role in aggression. However, over time, the focus shifted from examining the LS as a potential substrate for reward to how it mediates aggression and other social behaviors^8,9^. In particular, efforts were made to identify how the LS regulates aggressive behavior through its connectivity with the hypothalamus. One such pathway from the ventral LS (LSv) to the ventromedial hypothalamus (vmH) was found to inhibit aggression^10^. It was then determined that this inhibitory pathway is further regulated by dorsal LS (LSd) projections to the LSv^11^, suggesting that local microcircuits within the LS may serve to integrate information to shape behavioral outputs^12^. Additionally, social information was shown to be organized along the dorsoventral (D-V) axis of the LS, with LSd neurons responding more strongly to non-kin odors compared to kin-responsive LSv neurons^13^.

The dorsoventral organization of information is not restricted to social information within the LS. In fact, information about locomotion is organized along its various anatomical axes^14–17^. Place fields have also been identified in the LS, especially in the location of rewards^18,19^. However, the LS is not a homogenous structure and it contains many different subregions and cell types^20–22^. These cell types are organized in clusters along the D-V axis of the LS^23,24^. Furthermore, it has been hypothesized that the specific connectivity patterns of neurons in different regions along the D-V axis of the LS may contribute to different behavioral outcomes^12,16^. For example, one study demonstrated that molecularly distinct, spatially organized LS subpopulations receive biased hippocampal and hypothalamic inputs that determine whether they encode cue-or action-related information^25^. This may account for the functional heterogeneity that has been attributed to the LS, such as the high variability in the number of place cells across studies^26,27^.

Recently, there has been renewed interest in localizing the reward center within the LS, specifically due to its implication in substance use disorders^8,28^. One such study found that neurotensin-expressing LS neurons, but not somatostatin-expressing LS neurons, are activated during opioid withdrawal through increased glutamatergic input^29^. Another study identified a circuit by which glucagon-like peptide-1 receptor-expressing neurons in the LSd inhibited LSv neurons and modulated alcohol consumption through an interaction with the mesolimbic dopamine system^30^. Conversely, other efforts to localize reward signals have demonstrated that although stimulation of LSv neurons can drive place preference, their activity also correlates with stress-induced grooming and other states of heightened arousal^31^. Thus, it is still unclear how the LS promotes reward-seeking and how individual LS subregions may contribute to reward-related behaviors.

Here, we use cellular resolution calcium imaging to examine reward representations along the D-V axis of the LS. We find that there is a dorsoventral gradient of reward representations such that a greater proportion of LSd neurons are excited by reward and a greater proportion of LSv neurons are inhibited by reward. We also identify two LS neural populations that project to brain regions involved in context- and reward-encoding, the nucleus accumbens (NAc) and the bed nucleus of the stria terminalis (BNST), which occupy discrete dorsal and ventral compartments in the LS^32–34^. These two projection populations encode reward in a manner corresponding to the observed dorsoventral gradient. Consistent with this gradient, activation of the dorsal LS-NAc population and inhibition of the ventral LS-BNST population both support intracranial self-stimulation. Consequently, we have identified anatomically separate LS projection populations that have functionally distinct roles in reward-seeking behavior.

## Results

### Lateral septum neurons display a dorsoventral gradient of reward responses

To examine reward representations throughout the lateral septum (LS), we performed cellular resolution calcium imaging of LS neurons in freely behaving mice during a sucrose operant assay, in which mice are trained to nose-poke a choice port to access a 10 µL sucrose reward from a sucrose reward port on the opposite wall of the operant chamber (Figure 1A). Over several consecutive behavioral sessions, imaging mice completed a significantly greater number of sucrose trials and decreased their latency to consume the sucrose reward (Figure 1B,C; one-way ANOVA with post-hoc t tests, sucrose trials: p = 9.16*10^-9, reward latency: p = 2.31*10^-10, for post-hoc t tests see Supplementary Table 1). We imaged the calcium activity of individual LS neurons during each behavioral session by injecting an AAV expressing the calcium indicator GCaMP6f into the LS and implanting a gradient index (GRIN) lens at varying depths along the dorsoventral (D-V) axis of the LS in 15 mice (Figure 1D-F). CNMFe was then used to detect the activity of individual neurons and the fluorescence traces of each neuron were subsequently aligned to task events, such as choice port and sucrose reward port entry (Figure 1E)^35^. After all behavioral experiments, lens placement was reconstructed using Wholebrain software and imaging mice were divided into dorsal (LSd), intermediate (LSi), and ventral (LSv) groups depending on the D-V coordinate of their lens on post-hoc histology (Figure 1F,G)^36^.

**Figure 1.**
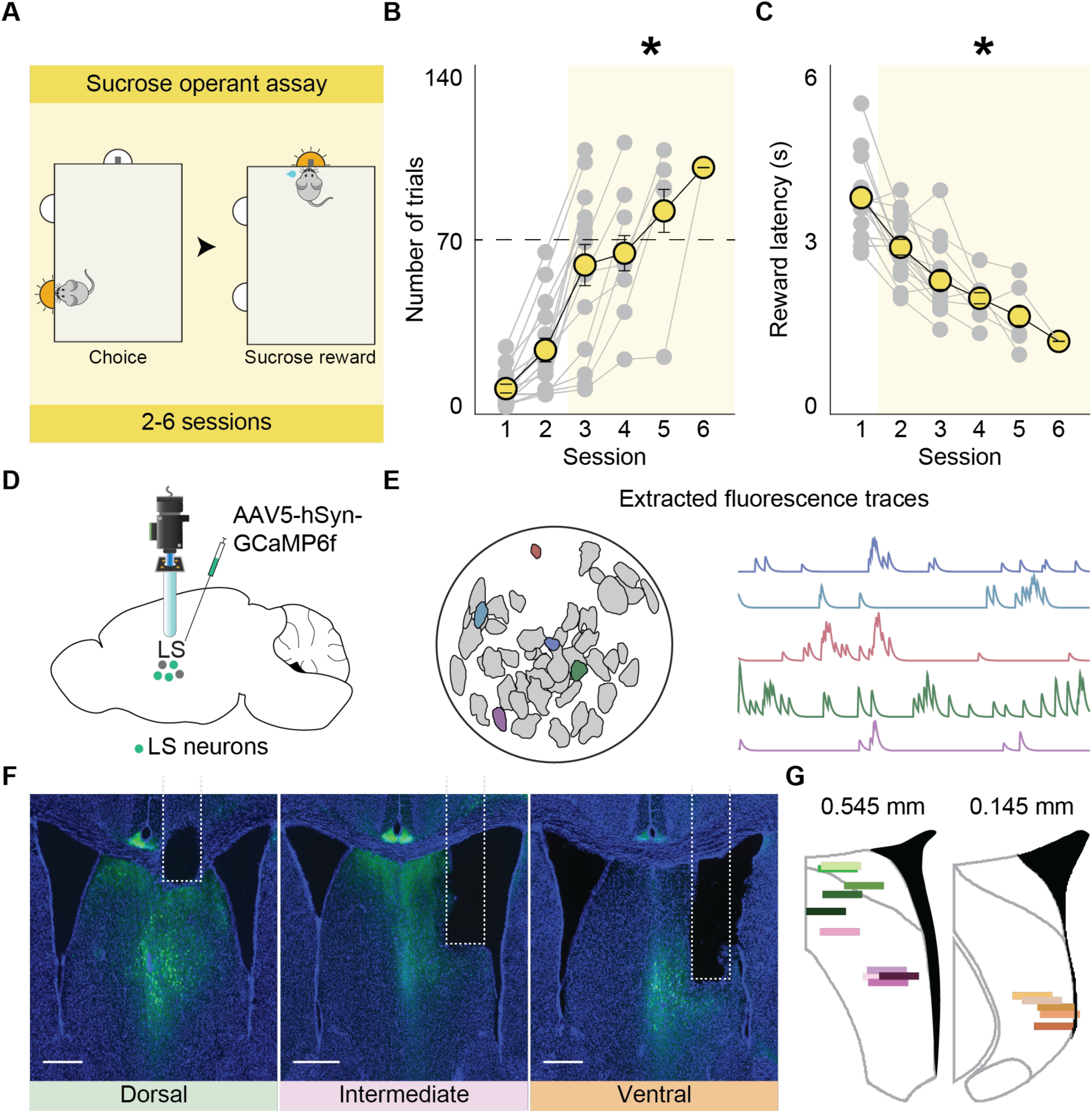
Cellular resolution calcium imaging along the D-V axis of the LS during a sucrose operant assay. A) Schematic of the sucrose operant assay. Mice are placed on restricted water access and trained to nose poke a choice port in order to access a sucrose reward in the sucrose reward port on the adjacent wall of the operant arena across consecutive sessions. B) With training on the sucrose operant assay, mice complete an increasing number of successful sucrose trials. The final session on the operant assay was determined as the session in which mice completed over 70 successful trials (indicated by the dashed black line). One-way ANOVA with post-hoc t tests (p = 9.16*10^-9). C) With training on the sucrose operant assay, mice show a significant decrease in latency to consume a sucrose reward (reward latency). One-way ANOVA with post-hoc t tests (p = 2.31*10^-10). For post-hoc t tests see Supplementary Table 1. Individual mice shown in grey, averages shown in yellow. N = 15 mice. Error bars indicate ± SEM. *p<0.05. D) Schematic of the viral strategy used to label LS neurons with GCaMP6f and the imaging setup of GRIN lens placement in the LS. The GRIN lens was placed at different D-V coordinates in individual mice to image from the dorsal, intermediate and ventral LS. E) Example field of view and extracted calcium traces corresponding to individual neurons. F) Representative images showing GRIN lens placement in different dorsal (left), intermediate (middle) and ventral (right) positions in the LS at 4x magnification. GCaMP6f expression shown in green and DAPI expression shown in blue. Lens tract outlined with white dashed lines. Scale bars represent 500 μm. G) Reconstruction of GRIN lens placement in dorsal (green, n = 5 mice), intermediate (pink, n = 5 mice) and ventral (orange, n = 5 mice) LS using Wholebrain software. Each colored line corresponds to an individual mouse and shows the position of the lens relative to bregma in the Allen Mouse Brain Common Coordinate Framework.

We quantified the number of neurons that were significantly modulated by choice port entry and sucrose reward consumption across all imaged neurons on the final session of the sucrose operant assay, which was determined to be the first session in which mice completed over 70 sucrose trials (Figure 1B, denoted by the dashed black line). We imaged a total of 931 neurons across 15 mice: 359 neurons from the LSd (Figure 1G, n = 5 mice, green), 303 neurons from the LSi (Figure 1G, n = 5 mice, pink), and 269 neurons from the LSv (Figure 1G, n = 5 mice, orange). Across dorsal, intermediate and ventral compartments of the LS, a greater proportion of neurons was modulated by reward consumption compared to choice port entry (Figure 2A: LSd: choice = 139/359 neurons, reward = 253/359 neurons; LSi: choice = 81/303 neurons, reward = 205/303 neurons; LSv: choice = 51/269 neurons, reward = 156/269 neurons; proportion z test, choice v reward in LSd, LSi and LSv: p <0.000001). Reward modulated neurons were further classified as either sucrose reward excite if they were positively modulated by reward (Figure 2B; Supplementary Figure 1A) or sucrose reward inhibit if they were negatively modulated by reward (Figure 2C; Supplementary Figure 1B) and the proportion of reward-responsive neurons was quantified along the D-V axis of the LS (Figure 2D-F).

**Figure 2.**
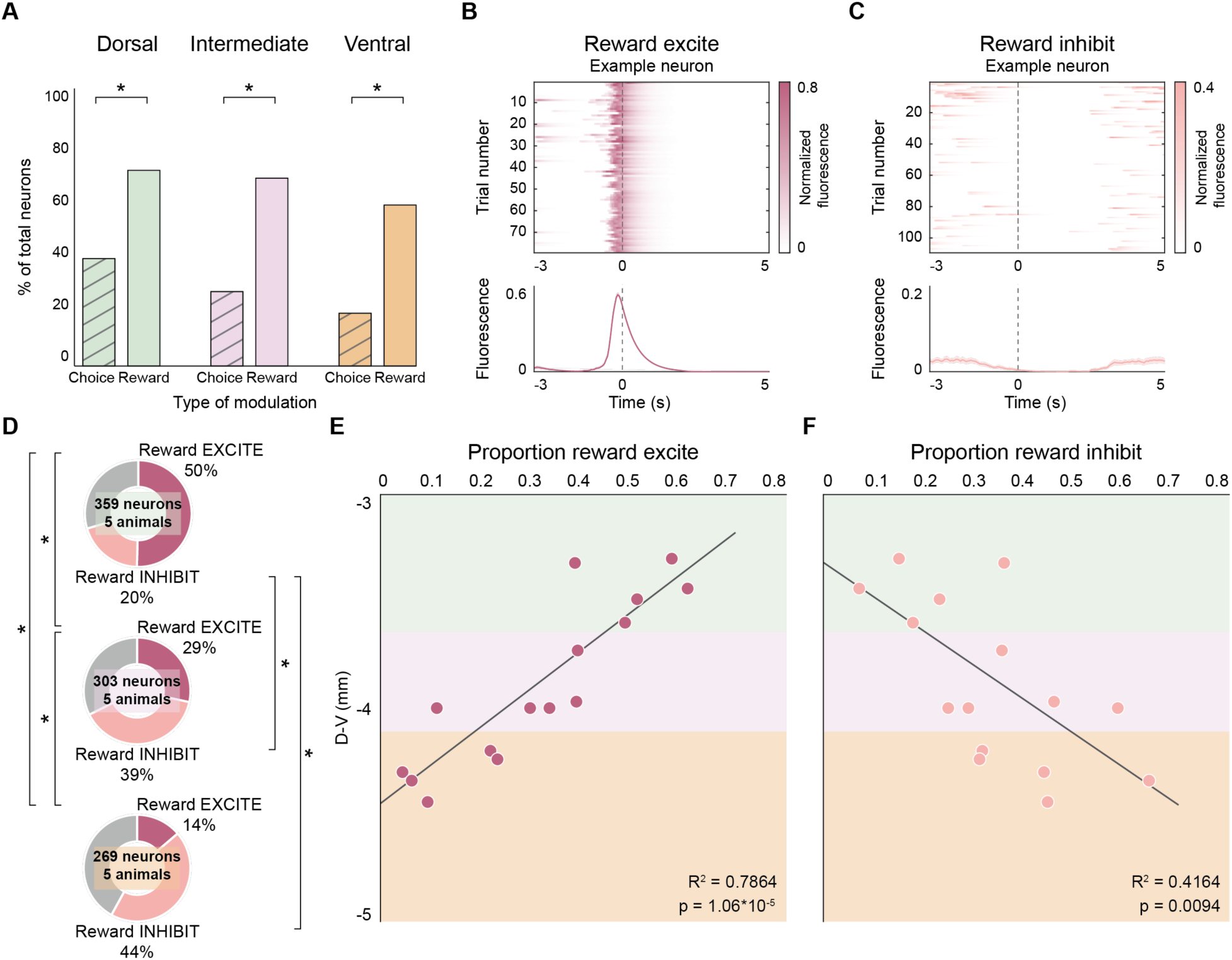
Proportion of sucrose reward responses forms a dorsoventral gradient in the LS. A) The proportion of neurons modulated by reward (LSd: n = 71%, 253/359 neurons; LSi: n = 68%, 205/303 neurons; LSv: 58%, 156/269 neurons) is greater than the proportion of those modulated by choice (LSd: n = 39%, 139/359 neurons; LSi: n = 27%, 81/303 neurons; LSv: 19%, 51/269 neurons) in all three compartments of the LS. Proportion z test (LSd, LSi, and LSv: p <0.000001). B) Heatmap of normalized fluorescence on each trial (top panel) and average fluorescence traces ± SEM (bottom panel) of an example neuron significantly positively modulated by sucrose reward (reward excite). C) Heatmap of normalized fluorescence on each trial (top panel) and average fluorescence traces ± SEM (bottom panel) of an example neuron significantly negatively modulated by sucrose reward (reward inhibit). Dashed line at zero indicates reward onset. D) Proportions of total recorded LS neurons that are reward excite or reward inhibit in the dorsal (top panel), intermediate (middle panel) and ventral (bottom panel) LS. A greater proportion of dorsal LS (LSd) neurons are excited by reward (n = 50%, 181/359 neurons), compared to those in the intermediate LS (LSi; n = 29%, 86/303) and ventral LS (LSv; n = 14%, 37/269). Proportion z test (LSd vs LSi: p = 8.52*10^-9; LSd vs LSv: p <0.000001; LSi vs LSv: p = 2.14*10^-5). Whereas, a greater proportion of LSv (n = 44%, 119/269 neurons) and LSi (n = 39%, 119/303 neurons) neurons are inhibited by reward compared to those in the LSd (n = 20%, 72/359). Proportion z test (LSd vs LSi: p = 5.41*10^-8; LSd vs LSv: p = 7.11*10^-11; LSi vs LSv: p = 0.229). E) The proportion of reward excite neurons imaged in individual mice is correlated to their D-V position in the LS (LSd highlighted in green, LSi highlighted in pink, and LSv highlighted in orange), such that the more dorsal the lens, the greater the proportion of reward excite neurons and the more ventral the lens, the smaller the proportion of reward excite neurons. Linear regression (p = 1.06*10^-5). F) The proportion of reward inhibit neurons imaged in individual mice is inversely correlated to their D-V position in the LS, such that the more ventral the lens, the greater the proportion of reward inhibit neurons and the more dorsal the lens, the smaller the proportion of reward inhibit neurons. Linear regression (p = 0.0094). *p<0.05. N = 15 mice, 5 mice per LS D-V compartment.

On the final day of behavior on the sucrose operant assay, across all LSd neurons, 50% (n = 181/359 neurons) were excited by sucrose reward and 20% (n = 72/359 neurons) were inhibited by sucrose reward (Figure 2D, top panel). Across all LSi neurons, 29% (n = 86/303 neurons) were excited by sucrose reward and 39% (n = 119/303) were inhibited by sucrose reward (Figure 2D, middle panel). Across all LSv neurons, 14% (n = 37/269) were excited by sucrose reward and 44% (n = 119/269) were inhibited by sucrose reward (Figure 2D, bottom panel). A significantly greater proportion of LSd neurons were excited by sucrose reward than LSi and LSv neurons and a greater proportion of LSi neurons were excited by sucrose reward than LSv neurons (Figure 2D; proportion z test, LSd v LSi: p = 8.52*10^-9, LSd v LSv: p <0.000001, LSi v LSv: p = 2.14*10^-5). Conversely, a significantly greater proportion of LSv and LSi neurons were inhibited by sucrose reward than LSd neurons (Figure 2D; proportion z test, LSd v LSi: p = 5.41*10^-8, LSd v LSv: 7.11*10^-11, LSi v LSv: p = 0.229).

Once we had established that the proportion of sucrose reward excite and inhibit neurons varied based on broad subregions within the LS, we next wanted to determine if lens position along the D-V axis was associated with the proportion of sucrose reward modulated neurons. To do this, we correlated these proportions for each individual mouse with the reconstructed D-V positions of the lens (Figure 2E,F). We found that the proportion of reward excite (Figure 2E; linear regression, p = 1.06*10^-5) and proportion of reward inhibit (Figure 2F; linear regression, p = 0.0094) neurons were significantly linearly correlated along the D-V axis of the LS, such that the more dorsal the lens, the greater the proportion of reward excite and the lesser the proportion of reward inhibit neurons, while the more ventral the lens, the lesser the proportion of reward excite and the greater the proportion of reward inhibit neurons.

### Anatomically and electrophysiologically distinct LS neurons project to the nucleus accumbens (NAc) and bed nucleus of the stria terminalis (BNST)

In previous work characterizing the anatomical organization of various projection populations in the LS, especially along its D-V axis, we showed that LS neurons projecting to the nucleus accumbens (LS-NAc) and to the bed nucleus of the stria terminalis (LS-BNST) occupied dorsal and ventral compartments of the LS, respectively^37^. To determine if there was any overlap between the two populations and to independently verify the localization of these two projections within the LS, we used a dual retrograde viral tracing strategy to label both LS-NAc and LS-BNST neurons within the same mouse (Figure 3A). We injected a retrogradely transported Cre virus and a retrogradely transported Flpo virus into either target population (NAc or BNST) and a Cre-dependent green fluorescent protein (GFP) and Flp-dependent virus labeled with an mCherry fluorophore into the LS (Figure 3A). This allowed us to label individual LS-NAc and LS-BNST neurons with either green or red fluorophores within the same mouse (Figure 3B). We then quantified the position of individual fluorescently labeled LS neurons by registering 50 µm coronal brain sections to the corresponding section from the Allen Brain Atlas using ABBA software^38^ at three different anteroposterior (A-P) coordinates (0.845, 0.545, and 0.02 mm anterior to bregma) to account for the anatomical extent of the LS, which spans ∼1 mm along the A-P axis of the brain^20^. Across all three A-P coordinates, a total of 3,057 fluorescently labeled neurons were detected (n = 6 mice, 3 mice per viral strategy). Only 0.8% (n = 24/3,057 neurons) were double-labeled, indicating minimal overlap between the two projection populations (Figure 3C). This separation of projection populations was consistent within individual mice (Figure 3D; LS-BNST: 97.6% ± 1.30% and LS-NAc: 99.1% ± 0.34% non-overlapping). Of the single-labeled neurons, significantly more belonged to the LS-NAc (n = 69.6%, 2,128/3,057 neurons) population than the LS-BNST (29.6%, 905/3,057) population (Figure 3C; proportion z test, p < 0.000001), corroborating prior evidence that the LS-NAc projection is denser than other LS projection populations^37^.

**Figure 3.**
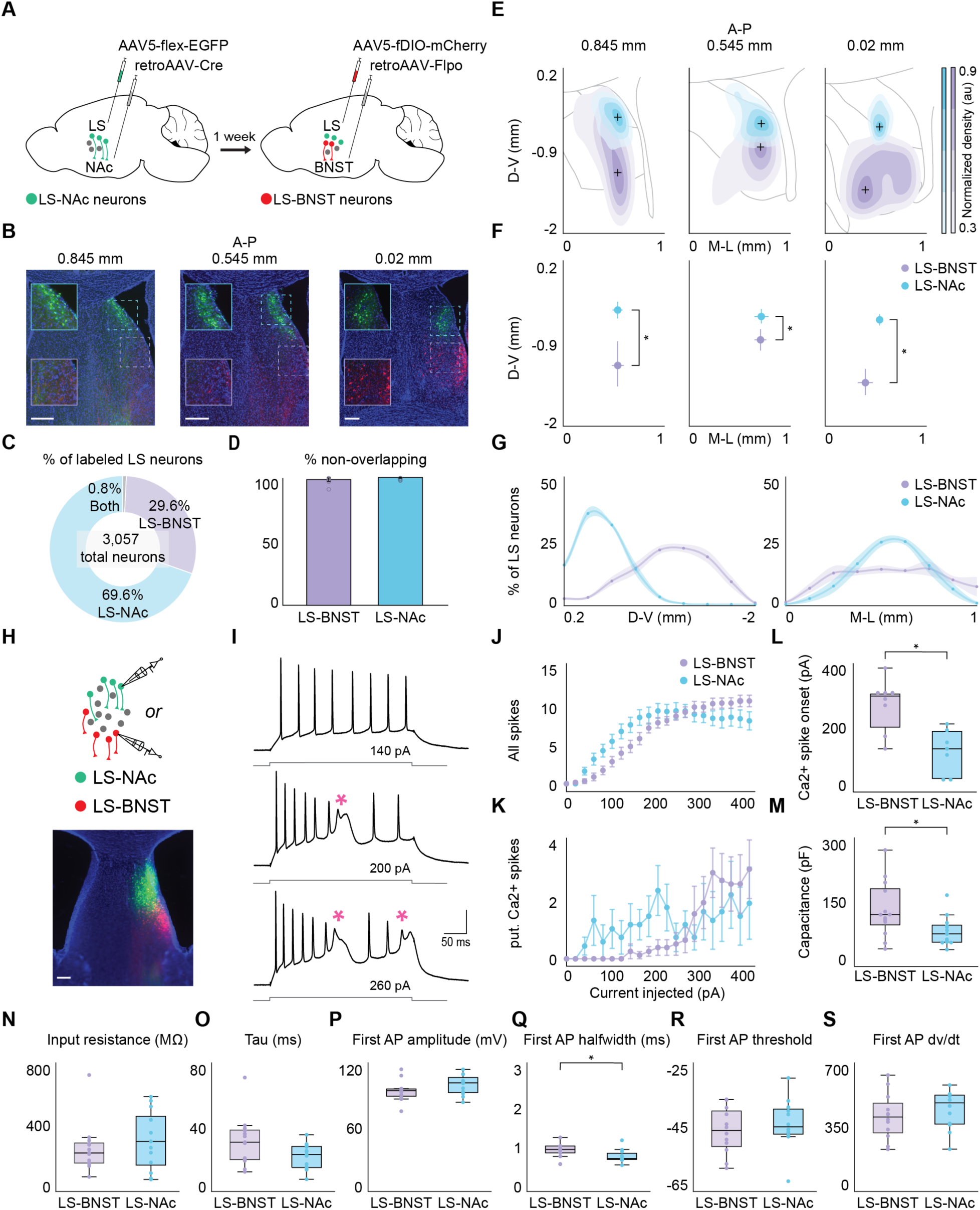
LS-NAc and LS-BNST neurons occupy distinct dorsal and ventral compartments of the LS and show differences in their electrophysiological properties. A) Schematic of the viral strategy used to label LS neurons that project to the nucleus accumbens (NAc) or the bed nucleus of the stria terminalis (BNST) within the same mouse. A retrogradely transported Cre virus is injected into the NAc and a Cre-dependent GFP is injected into the LS. In addition, a retrogradely transported Flpo virus is injected into the BNST and a Flp-dependent mCherry in the LS. B) Representative images of LS-NAc (GFP) and LS-BNST (mCherry) neurons at 3 different A-P coordinates at 4x magnification. Insets show areas outlined by blue (LS-NAc, GFP) or purple (LS-BNST, mCherry) squares at higher magnification. GFP expression shown in green, mCherry expression shown in red and DAPI expression shown in blue. Scale bars represent 100 μm. C) Proportion of total counted neurons that were labeled with both fluorophores (grey, both: n = 0.8%, 24/3,057 neurons), the fluorophore corresponding to the LS-BNST population (purple, LS-BNST: n = 29.6%, 905/3,057), and the fluorophore corresponding to the LS-NAc population (blue, LS-NAc: n =69.6%, 2,128/3,057 neurons). D) LS-BNST (purple) and LS-NAc (blue) neurons are largely non-overlapping with each other. Individual colored points correspond to individual mice. Error bars indicate ± SEM. E) Density contour plots showing the location of LS-NAc (blue) and LS-BNST (purple) projection neurons along the D-V and mediolateral (M-L) axes (relative to the corpus callosum) across all mice at 3 A-P coordinates (0.845 mm, 0.545 mm, and 0.02 mm, from left to right). Each contour corresponds to a normalized density of 30%, 50%, 70%, and 90% (from lightest to darkest shade of the color representing each projection population). The gray outline represents the anatomical outline of the LS at that A-P coordinate, including a partial outline of the lateral ventricle. F) LS-NAc neurons (blue) are clustered in the dorsal LS, while LS-BNST (purple) neurons are clustered in the ventral LS across various A-P coordinates, as shown by the centroids of the 90% contour plots with the horizontal and vertical spread of the 90% contour at 3 A-P coordinates (0.845 mm, 0.545 mm, and 0.02 mm, from left to right). The distribution of LS-NAc and LS-BNST neurons are separable at all A-P coordinates. Pairwise KS tests (at the A-P coordinate of 0.845 mm: p = 7.66*10^-60, at the A-P coordinate of 0.545 mm: p = 5.76*10^-54, at the A-P coordinate of 0.02 mm: p = 5.29*10^-43). G) D-V (left panel) and M-L (right panel) distributions of proportions of LS-NAc (blue) and LS-BNST (purple) neurons in the LS relative to the corpus callosum. Shaded error regions indicate ± SEM. N = 6 mice. H) Using the same retrograde viral tracing strategy, LS-NAc and LS-BNST neurons were labeled with different fluorophores and targeted for electrophysiological recordings (top panel). A representative image of a brain section that was used for electrophysiological recordings with LS-NAc (GFP) and LS-BNST (mCherry) neurons at 4x magnification (bottom panel). GFP expression shown in green, mCherry expression shown in red and DAPI expression shown in blue. Scale bar represents 100 μm. I) Example voltage traces recorded from LS neurons at current injections of varying intensities (top: 140 pA, middle: 200 pA, bottom: 260 pA). Putative calcium spikes denoted by pink asterisks. J) LS-BNST (purple, n = 13 neurons) and LS-NAc (blue, n = 13 neurons) neurons generate a similar number of sodium spikes at various injected currents (pA). Sidak’s multiple comparisons tests. K) LS-BNST (purple) and LS-NAc (blue) neurons generate a similar number of putative calcium (Ca2+) spikes at various injected currents (pA). Sidak’s multiple comparisons tests. See Supplementary Table 1. Error bars indicate ± SEM. L) Of the recorded neurons that generate calcium spikes, LS-NAc neurons (n = 7 neurons) produce calcium (Ca2+) spikes at a lower injected current (pA) than LS-BNST neurons (n = 8 neurons). Unpaired t test (p = 0.0025). M) Recorded LS-NAc neurons (blue, n = 13 neurons) have a lower capacitance than LS-BNST neurons (purple, n = 13 neurons). Unpaired t test (p = 0.021). N) There was no difference in the input resistance of LS-NAc (blue) and LS-BNST (purple) neurons. Unpaired t test (p = 0.407). O) There was no difference in the membrane time constant of LS-NAc (blue) and LS-BNST (purple) neurons. Unpaired t test (p = 0.0619). P) The amplitudes of the first evoked sodium spike were comparable between LS-NAc (blue, n = 12 neurons) and LS-BNST (purple, n = 12 neurons) neurons. Unpaired t test (p = 0.178). Q) The halfwidth of the first evoked sodium spike of LS-BNST (purple) neurons was greater than that of LS-NAc (blue) neurons. Unpaired t test (p = 0.0313). R) The thresholds of the first evoked sodium spike were similar between LS-NAc (blue) and LS-BNST (purple) neurons. Unpaired t test (p = 0.464). S) The first evoked sodium spike dv/dt were comparable between LS-NAc (blue) and LS-BNST (purple) neurons. Unpaired t test (p = 0.457). *p<0.05. Boxplots: center line denotes median, box edges indicate the lower (25th) and upper (75th) quartiles, and whiskers extend to ±1.5*IQR. Individual colored points correspond to individual recorded neurons.

Prior work of ours also demonstrated that different LS projection populations occupy discrete regions of the LS^37^. Therefore, to compare the distribution of LS-NAc and LS-BNST populations along the anatomical extent of the LS within individual mice, we created density contour plots of the two projection populations at 3 separate A-P coordinates (relative to bregma) that cover the anterior (0.845 mm), intermediate (0.545 mm), and posterior (0.02 mm) LS. At each A-P coordinate, the D-V and mediolateral (M-L) distribution (relative to the center of the corpus callosum) of LS projection neurons, collapsed across 6 mice, was determined (Figure 3E). For all 3 A-P coordinates, the 90% contours of the LS-NAc and LS-BNST distributions were significantly different from each other (Figure 3F; pairwise KS tests, A-P coordinate of 0.845 mm: p = 7.66*10^-60, A-P coordinate of 0.545 mm: p = 5.76*10^-54, and A-P coordinate of 0.02 mm: p = 5.29*10^-43). By plotting the D-V and M-L coordinate of the centroid of the density contour plots (Figure 3F), we found that the LS-NAc projection population consistently resides more dorsally in the LS (LS-NAc centroid D-V coordinate from anterior to posterior: -0.455, -0.541, and -0.584) when compared to the LS-BNST projection population (LS-BNST centroid D-V coordinate from anterior to posterior: -1.19, -0.851, and -1.42), but that the two are relatively evenly distributed along the M-L axis of the LS, except in the most posterior portion of the LS (LS-NAc centroid M-L coordinate from anterior to posterior: 0.540, 0.713, and 0.538; LS-BNST centroid M-L coordinate from anterior to posterior: 0.540, 0.706, and 0.397). This anatomical organization of LS projection population neurons can also be visualized through the D-V and M-L distributions of the proportions of LS neurons of each projection population (Figure 3G).

Next, to determine if these anatomically distinct populations also differed in their electrophysiological properties, we performed fluorescence guided whole cell recordings of LS-NAc and LS-BNST neurons using the aforementioned dual retrograde viral strategy (Figure 3H, n = 3 mice, LS-BNST: n = 13 neurons, LS-NAc: n = 13 neurons). We found that both populations produced sodium spikes, as well as putative Ca2+ spikes^39^ in response to current injections of various intensities (Figure 3I-K). We found that LS-NAc neurons produced putative Ca2+ spikes at lower current intensities compared to LS-BNST neurons (Figure 3L; unpaired t test, p = 0.00250). LS-BNST neurons showed greater whole-cell capacitance than LS-NAc neurons (Figure 3M; unpaired t test, p = 0.0210). We found no differences in the input resistance or membrane time constant between both types of neurons (Figure 3N,O; unpaired t test, input resistance: p = 0.407, tau: p = 0.0619). Additionally, active properties of the first evoked sodium spike were comparable between the two populations except for the spike halfwidth (Figure 3P-S; unpaired t test, amplitude: p = 0.178, halfwidth: p = 0.0313, threshold: p = 0.464, dv/dt: p = 0.457). Since spike amplitude, threshold and dv/dt did not differ, the greater halfwidth of LS-BNST spikes likely reflects differences in repolarizing conductances rather than in the core sodium spike–generating machinery, which appears to be broadly shared by the two projection populations. Together, these findings show that LS-NAc and LS-BNST neurons differ in their intrinsic excitability and passive membrane properties while sharing core spike-generating machinery, suggesting that these two populations may serve distinct roles in reward processing.

### LS projection populations encode reward differently

To examine whether the dorsal LS-NAc and ventral LS-BNST neurons differently process reward-related information and contribute to the observed dorsoventral gradient of reward representations in our non-specific LS imaging, we imaged the activity of individual LS-NAc (n = 407 neurons, 6 mice) and LS-BNST (n = 121 neurons, 6 mice) neurons during the sucrose operant assay (Figure 4A). In order to target individual LS projection neurons, we injected a retrogradely transported Cre virus labeled with mCherry into either the NAc or BNST and a Cre-dependent GCaMP6f into the LS and implanted a GRIN lens over the individual projection population (Figure 4A-C). We then trained mice on the same sucrose operant assay (Figure 1A). Mice acquired the task over several consecutive sessions. As with the non-specific imaging cohort (Figure 1B,C), with training, all projection-specific imaging mice increased the number of sucrose trials and decreased their latency to consume the sucrose reward (Figure 4D,E; one-way ANOVA with post-hoc t tests, sucrose trials: p = 2.50*10^-4, reward latency: p = 0.012, for post-hoc t tests see Supplementary Table 1). We then quantified the reward responses on the final behavioral session, which was the session in which mice completed more than 70 sucrose trials (Figure 4D, denoted by the dashed black line). A greater proportion, 62% (n = 254/407 neurons), of all imaged LS-NAc neurons were excited by sucrose reward compared to 6% (n = 7/121 neurons) of LS-BNST neurons (Figure 4F; proportion z test, p <0.000001), while a greater proportion, 59% (n = 72/121 neurons), of LS-BNST neurons were inhibited by sucrose reward than the 15% (n = 60/407 neurons) of LS-NAc neurons (Figure 4F; proportion z test, p <0.000001). We also determined that the D-V coordinate of the lens correlated with the proportion of reward excite and reward inhibit neurons identified in individual mice. The dorsally clustered LS-NAc imaging mice had a greater proportion of reward excite neurons than the ventrally clustered LS-BNST imaging mice (Figure 4H; linear regression, p = 6.42*10^-6), while the LS-BNST imaging mice had a greater proportion of reward inhibit neurons than the LS-NAc imaging mice (Figure 4I; linear regression, p = 6.09*10^-5). Furthermore, we could predict the identity of the projection population with higher accuracy than shuffled data using decoders trained on the neural responses in a 6 second window around reward consumption (Figure 4G; unpaired t test, p = 6.57*10^-53, see *Decoding projection population using neural data* in the Methods section).

**Figure 4.**
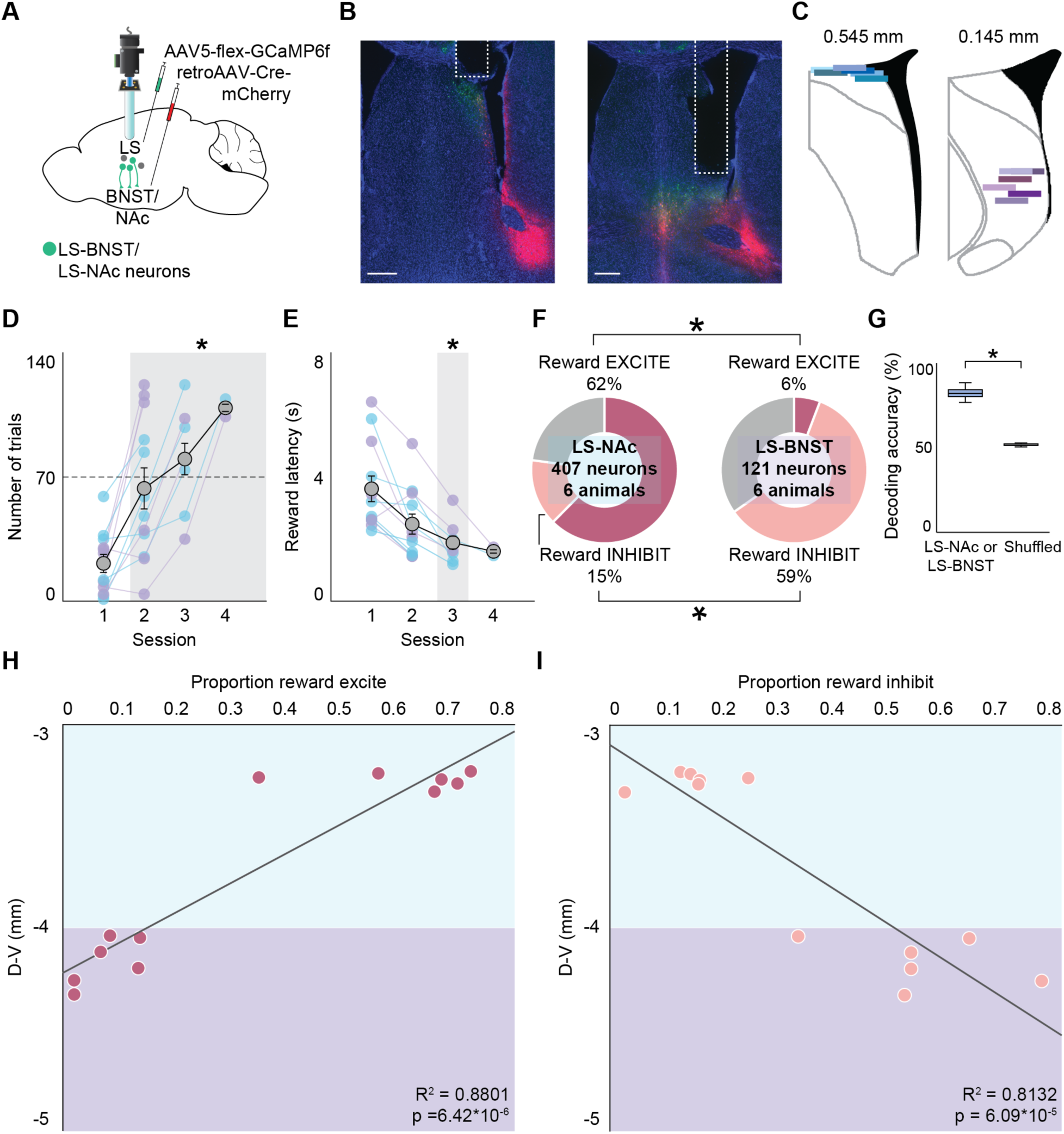
Differential reward representations in LS projection populations located along its D-V axis. A) Schematic of the viral strategy used to label LS neurons that project to either the NAc or BNST with GCaMP6f and the target brain region with mCherry, along with the imaging setup of GRIN lens placement in the LS. The GRIN lens was positioned more dorsally to image LS-NAc neurons and more ventrally to image LS-BNST neurons. B) Representative images showing GRIN lens placement for LS-NAc (left) and LS-BNST (right) projection populations at 4x magnification. GCaMP6f expression shown in green, mCherry expression in the target brain region shown in red and DAPI expression shown in blue. Tract from lens outlined in white. Scale bars represent 500 μm. C) Reconstruction of GRIN lens placement in LS-NAc (blue, n = 6 mice) and LS-BNST (purple, n = 6 mice) imaging mice using Wholebrain software. Each colored line corresponds to an individual mouse and shows the position of the lens relative to bregma in the Allen Mouse Brain Common Coordinate Framework. D) With training on the sucrose operant assay, mice complete an increasing number of successful sucrose trials. One-way ANOVA with post-hoc t tests (p = 2.50*10^-4). E) With training on the sucrose operant assay, mice show a significant decrease in latency to consume a sucrose reward. One-way ANOVA with post-hoc t tests (p = 0.012). For post-hoc t tests see Supplementary Table 1. Individual mice shown in blue (indicates LS-NAc, n = 6 mice) and purple (indicates LS-BNST, n = 6 mice), averages shown in yellow. Error bars indicate ± SEM. *p<0.05. F) A greater proportion of LS-NAc neurons are excited by sucrose reward (n = 62%, 254/407 neurons) compared to LS-BNST (n = 6%, 7/121 neurons). Proportion z test (p < 0.000001). In contrast, a greater proportion of LS-BNST neurons are inhibited by sucrose reward (n = 59%, 72/121) compared to LS-NAc (15%, 60/407 neurons). Proportion z test (p < 0.000001). G) Decoders trained on the neural data in a 6s window around the reward are able to decode the identity of the projection population with higher accuracy than a shuffled dataset. Unpaired t test (p = 6.57*10^-53). H) The proportion of reward excite neurons imaged in LS projection populations in individual mice is correlated to the D-V position of the lens (LS-NAc highlighted in blue and LS-BNST highlighted in purple), such that the more dorsal the lens, the greater the proportion of reward excite neurons and the more ventral the lens, the smaller the proportion of reward excite neurons. Linear regression (p = 6.42*10^-6). I) The proportion of reward inhibit neurons imaged in LS projection populations in individual mice is correlated to the D-V position of the lens, such that the more ventral the lens, the greater the proportion of reward inhibit neurons and the more dorsal the lens, the smaller the proportion of reward inhibit neurons. Linear regression (p = 6.09*10^-5).

To confirm that the neural responses we observed on the sucrose operant assay were specific to reward consumption and not broadly related to salient stimuli, we recorded LS projection population neurons on an assay in which mice were exposed to an aversive airpuff stimulus and on an additional session of the sucrose operant assay (Supplementary Figure 2A). We tracked individual neurons (Supplementary Figure 2B,C) and compared neural responses (Supplementary Figure 2D,E) across all three behavioral sessions. In LS-NAc mice, we recorded significantly fewer neurons in the airpuff session relative to the two operant assay sessions (Supplementary Figure 2C; one-way ANOVA with post-hoc t tests, p = 1.60*10^-4). In contrast, in the LS-BNST mice, we recorded a similar number of neurons across all sessions (Supplementary Figure 2C; one-way ANOVA, p = 0.846). Despite this discrepancy in number of recorded neurons, we were able to track a substantial number of individual neurons across sessions using CellReg software^40^ and examine their response profile to sucrose reward and airpuff (Supplementary Figure 2D,E). In both projection populations, we identified a subset of neurons that were significantly modulated by airpuff, though given the difference in number of recorded LS-NAc neurons between behavioral sessions, this proportion was less in LS-NAc mice, but not LS-BNST mice, when normalized to the number of neurons recorded in the second operant assay session (Supplementary Figure 2F; paired t test, LS-NAc: p = 0.0195, LS-BNST: p = 0.738). We then compared the degree of overlap between neurons that were modulated by sucrose reward in the operant assay sessions and those that were modulated by airpuff in the aversive exposure session. We found that there was significantly less overlap between the airpuff and sucrose operant assay conditions compared to the amount of overlap between the two operant assay sessions in LS-NAc mice, with similar findings in LS-BNST mice (Supplementary Figure 2G; one-way ANOVA with post-hoc t tests, LS-NAc: p = 1.31*10^-4, LS-BNST: p = 0.00546, for post-hoc t tests see Supplementary Table 1). As a result, reward-responsive LS projection population neurons appear to have relatively stable reward representations across operant assay sessions and are largely non-overlapping with the neurons that respond to aversive stimuli.

### Activation of LS-NAc neurons and inhibition of LS-BNST neurons independently elicits intracranial self-stimulation

Although historically the LS has been shown to robustly promote intracranial self-stimulation (ICSS), the substrate within the LS that mediates such positive reinforcement has remained elusive. Given the differences that we observed in reward representations across the two LS projection populations, we next asked whether optogenetic activation of these populations supports ICSS. We labeled LS-NAc or LS-BNST neurons with an excitatory opsin (channelrhodopsin, ChR2) or GFP (control) by bilaterally injecting a retrogradely transported Cre virus labeled with mCherry into the NAc or BNST and a Cre-dependent ChR2 (experimental, LS-NAc: n = 6 mice, LS-BNST: n = 5 mice) or GFP (control, n = 6 mice, 3 mice per projection population) into the LS, followed by the implantation of optical ferrules bilaterally into the LS (Figure 5A,B). Mice were then tested on an optogenetic ICSS paradigm, whereby nose-poking one choice port delivered 2s of 15 Hz blue light stimulation (Figure 5C, stim trial) and nose-poking the other choice port delivered no stimulation (Figure 5C, null trial). We compared the number of stimulated and null trials completed by experimental and control mice over three consecutive hour long sessions. We found that activation of LS-NAc, but not LS-BNST, neurons promoted ICSS compared to controls over three sessions (Figure 5D,E). Specifically, LS-NAc mice completed a significantly greater number of stimulated trials on the second and third sessions than LS-BNST and control mice (Figure 5D; two-factor ANOVA with group type (LS-NAc ChR2/LS-BNST ChR2/GFP) and session number (1/2/3) as factors, interaction: p = 2.99*10^-4, group type: p = 3.30*10^-10, session number: p = 0.446 with post-hoc unpaired t tests comparing group type within session type, session 1 - LS-NAc v control: p = 0.984, LS-BNST v control: p = 1, LS-NAc v LS-BNST: p = 1; session 2 - LS-NAc v control: p = 2.96*10^-5, LS-BNST v control: p = 1, LS-NAc v LS-BNST: p = 2.03*10^-4; session 3 - LS-NAc v control: p = 3.81*10^-7, LS-BNST v control: p = 1, LS-NAc v LS-BNST: p = 3.91*10^-6). Experimental and control mice did not differ in the number of null trials completed within sessions (Figure 5E; two-factor ANOVA with group type (LS-NAc ChR2/LS-BNST ChR2/GFP) and session number (1/2/3) as factors, interaction: p = 0.483, group type: p = 0.017, session number: p = 8.58*10^-4), the main effect of session number was driven by a decrease in the number of null trials completed over time (for post-hoc t tests see Supplementary Table 1).

**Figure 5.**
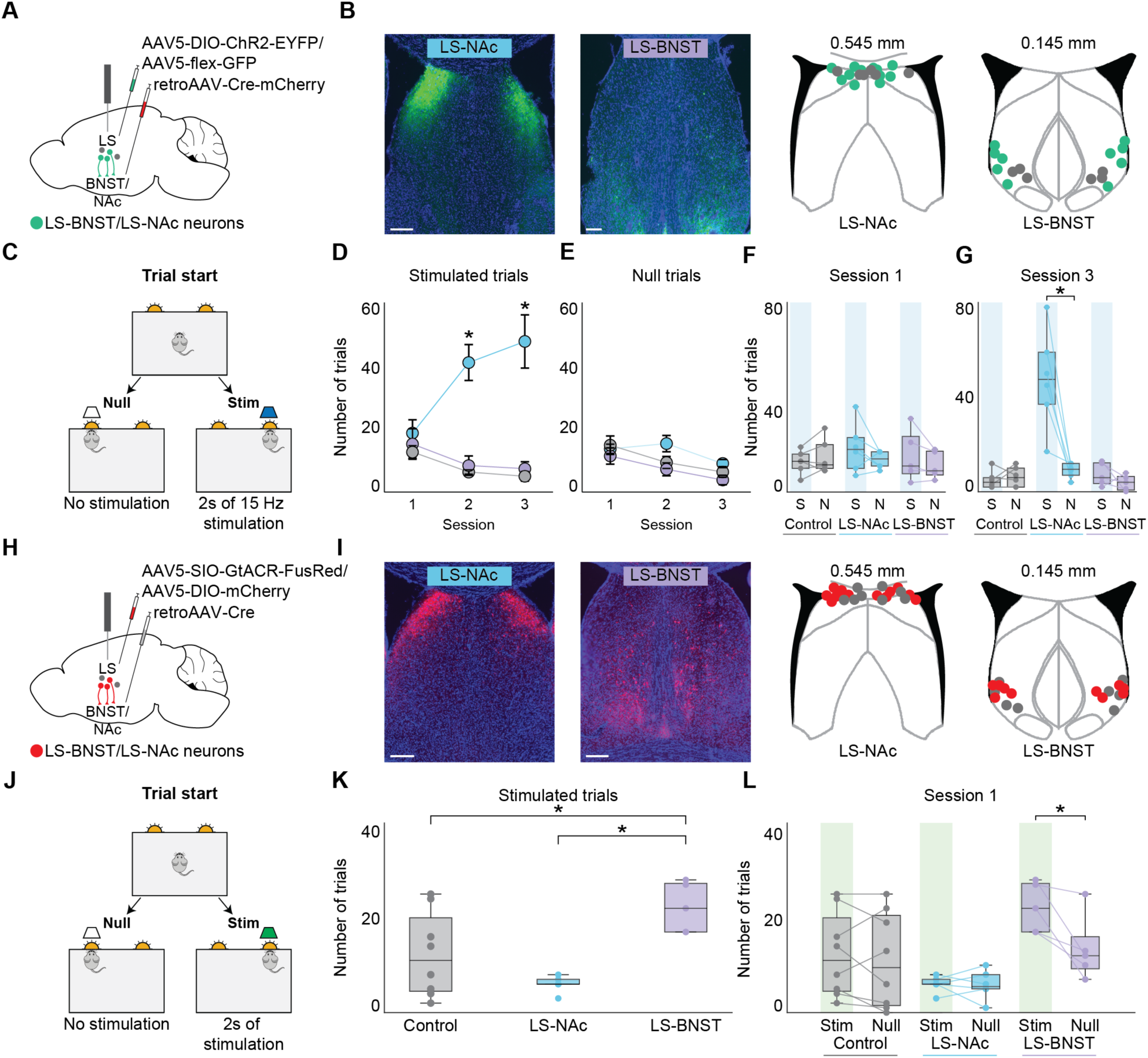
Opposing types of optogenetic manipulation of LS projection neurons produce intracranial self-stimulation (ICSS). A) Schematic of the viral strategy used to label LS-NAc or LS-BNST neurons with channelrhodopsin (experimental, LS-NAc: n = 6 mice, LS-BNST: n = 5 mice) or GFP (control, n = 6 mice, 3 mice per projection population) and the target brain region (BNST or NAc) with mCherry, along with optic ferrule placement in the LS. B) Representative images showing bilateral LS-NAc (left) and LS-BNST (right) channelrhodopsin labeling in the LS at 4x magnification, followed by reconstruction of optic ferrule placement. Channelrhodopsin (ChR2) expression shown in green and DAPI expression shown in blue. Scale bars represent 250 μm. Each colored dot in the reconstruction corresponds to a ferrule in an individual mouse (green are experimental mice and grey are control mice) and shows the position of the ferrule relative to bregma in the Allen Mouse Brain Common Coordinate Framework. C) Optogenetic ICSS behavioral paradigm. In a behavioral arena, mice can nose-poke two choice ports over the course of a one hour session. One choice port corresponds to 2 seconds of 15 Hz blue light stimulation (right panel, stimulated trial) and the other choice port corresponds to no optical stimulation (left panel, null trial). D) LS-NAc mice (blue, n = 6 mice) complete significantly more self-stimulation trials over three consecutive ICSS behavioral sessions compared to both LS-BNST (purple, n = 5 mice) and control mice (grey, n = 6 mice). Two-factor ANOVA with virus (LS-NAc ChR2/LS-BNST ChR2/GFP) and session number (1/2/3) as factors (interaction: p = 2.99*10^-4, virus: p = 3.30*10^-10, session number: p = 0.446 with post-hoc unpaired t tests comparing virus within session type, session 1 - LS-NAc v control: p = 0.984, LS-BNST v control: p = 1, LS-NAc v LS-BNST: p = 1; session 2 - LS-NAc v control: p = 2.96*10^-5, LS-BNST v control: p = 1, LS-NAc v LS-BNST: p = 2.03*10^-4; session 3 - LS-NAc v control: p = 3.81*10^-7, LS-BNST v control: p = 1, LS-NAc v LS-BNST: p = 3.91*10^-6). E) Experimental LS-NAc and LS-BNST mice complete a similar number of trials without optical stimulation as control mice during each ICSS behavioral session with an overall decrease in number of trials over three consecutive sessions. Two-factor ANOVA with virus (LS-NAc ChR2/LS-BNST ChR2/GFP) and session number (1/2/3) as factors (interaction: p = 0.483, virus: p = 0.0167, session number: p = 8.58*10^-4). F) During the first behavioral session, control (grey), LS-NAc (blue), and LS-BNST (purple) mice complete a similar number of stimulated and null trials. Mixed repeated measures ANOVA with virus (LS-NAc ChR2/LS-BNST ChR2/GFP) as the between subjects variable and trial type (stimulated/null) as the within subjects variable (interaction: p = 0.176, trial type: p = 0.193). G) During the third behavioral session, LS-NAc (blue) mice complete more stimulated trials than null trials, while control (grey) and LS-BNST (purple) mice complete a similar number of stimulated and null trials. Mixed repeated measures ANOVA with virus (LS-NAc ChR2/LS-BNST ChR2/GFP) as the between subjects variable and trial type (stimulated/null) as the within subjects variable (interaction: p = 2.67*10^-4, trial type: p = 9.03*10^-4 with post-hoc t tests comparing trial type within virus type, control: p = 0.811, LS-BNST: p = 0.557, LS-NAc: p = 5.46*10^-6). N = 17 mice, LS-NAc: n = 6 mice, LS-BNST: n = 5 mice, control: n = 6 mice. H) Schematic of the viral strategy used to label LS-NAc or LS-BNST neurons with the inhibitory opsin, GtACR (experimental, LS-NAc: n = 6 mice, LS-BNST: n = 5 mice) or mCherry (control, n = 8 mice, 4 mice per projection population) along with optic ferrule placement in the LS. I) Representative image showing bilateral LS-NAc (left) and LS-BNST (right) GtACR labeling in the LS at 4x magnification, followed by reconstruction of optic ferrule placement. GtACR expression shown in red and DAPI expression shown in blue. Scale bars represent 250 μm. Each colored dot in the reconstruction corresponds to a ferrule in an individual mouse (red are experimental mice and grey are control mice) and shows the position of the ferrule relative to bregma in the Allen Mouse Brain Common Coordinate Framework. J) Optogenetic ICSS behavioral paradigm for inhibition of LS-NAc or LS-BNST neurons. Mice can nose-poke two choice ports over the course of a one hour session. One choice port is associated with 2 seconds of continuous green light stimulation (right panel, stimulated trial) and the other choice port corresponds to no optical stimulation (left panel, null trial). K) During an ICSS behavioral session, LS-BNST (purple) mice complete a greater number of stimulated trials than LS-NAc (blue) and control (grey) mice. One-way ANOVA (p = 0.00319 with post-hoc t tests comparing between virus type, control v LS-NAc: p = 0.212, LS-BNST v control: p = 0.0425, LS-NAc v LS-BNST: p = 0.00235). L) During an ICSS behavioral session, LS-BNST (purple) mice complete more stimulated trials than null trials, while control (grey) and LS-NAc (blue) mice complete a similar number of stimulated and null trials. Mixed repeated measures ANOVA with virus (LS-NAc GtACR/LS-BNST GtACR/mCherry) as the between subjects variable and trial type (stimulated/null) as the within subjects variable (interaction: p = 0.00846, trial type: p = 0.0193 with post-hoc t tests comparing trial type within virus type, control: p = 0.477, LS-NAc: p = 0.934, LS-BNST: p = 9.23*10^-4). N = 19 mice, LS-NAc: n = 6 mice, LS-BNST: n = 5 mice, control: n = 8 mice. *p<0.05. Error bars indicate ± SEM. Boxplots: center line denotes median, box edges indicate the lower (25th) and upper (75th) quartiles, and whiskers extend to ±1.5*IQR. Individual colored points correspond to individual mice. For post-hoc t tests see Supplementary Table 1.

On the first session of the ICSS paradigm, all mice completed a similar number of stimulated and null trials (Figure 5F; mixed repeated measures ANOVA with group type (LS-NAc ChR2/LS-BNST ChR2/GFP) as the between subjects variable and trial type (stimulated/null) as the within subjects variable, interaction: p = 0.176, trial type: p = 0.193). However, by the third session on the ICSS paradigm, LS-NAc, but not LS-BNST or control mice, completed significantly more stimulated than null trials (Figure 5G; mixed repeated measures ANOVA with group type (LS-NAc ChR2/LS-BNST ChR2/GFP) as the between subjects variable and trial type (stimulated/null) as the within subjects variable, interaction: p = 2.67*10^-4, trial type: p = 9.03*10^-4 with post-hoc t tests comparing trial type within virus group, control: p = 0.811, LS-NAc: p = 5.46*10^-6, LS-BNST: p = 0.557).

Control mice that corresponded to either LS-NAc or LS-BNST projection populations (Supplementary Figure 3A) did not differ in number of stimulated (Supplementary Figure 3B; two-factor ANOVA with group type (LS-NAc controls/LS-BNST controls) and session number (1/2/3) as factors, interaction: p = 0.543, group type: p = 0.0791, session number: p = 0.0139) or null (Supplementary Figure 3C; two-factor ANOVA with group type (LS-NAc controls/LS-BNST controls) and session number (1/2/3) as factors, interaction: p = 0.806, group type: p = 0.519, session number: p = 0.0118) trials that they completed, so they were combined into one control group based on light stimulation protocol.

We then investigated how optogenetic inhibition of LS projection neurons affects reinforcement on the ICSS paradigm. We labeled LS-NAc or LS-BNST neurons with an inhibitory opsin (GtACR) or mCherry fluorophore (control) by bilaterally injecting a retrogradely transported Cre virus into the NAc or BNST and a Cre-dependent GtACR (experimental, LS-NAc: n = 6 mice, LS-BNST: n = 5 mice) or mCherry (control, n = 8 mice, 4 mice per projection population) into the LS, followed by the implantation of optical ferrules bilaterally into the LS (Figure 5H,I). We allowed mice to perform on the optogenetic ICSS paradigm whereby nose-poking one choice port resulted in 2s of sustained green light stimulation (Figure 5J, stim trial) and nose-poking the other choice port resulted in no stimulation (Figure 5J, null trial). On a single session of the ICSS paradigm, LS-BNST mice completed a greater number of stimulated trials compared to controls and LS-NAc mice (Figure 5K; one-way ANOVA, p = 0.00319 with post-hoc unpaired t tests comparing between group type, LS-NAc v control: p = 0.212, LS-BNST v control: p = 0.0425, LS-NAc v LS-BNST: p = 0.00235). Additionally, LS-BNST mice completed more stimulated than null trials, while control and LS-NAc mice completed a similar number of both trial types (Figure 5L; mixed repeated measures ANOVA with group type (LS-NAc GtACR/LS-BNST GtACR/mCherry) as the between subjects variable and trial type (stimulated/null) as the within subjects variable, interaction: p = 0.00846, trial type: p = 0.0193 with post-hoc t tests comparing trial type within virus group, control: p = 0.477, LS-NAc: p = 0.934, LS-BNST: p = 9.23*10^-4). Overall, the findings from these optogenetic manipulation experiments suggest that both optogenetic activation of LS-NAc neurons and optogenetic inhibition of LS-BNST neurons promote positive reinforcement on the ICSS paradigm, though on different time scales.

### Optogenetic activation of LS-NAc neurons promotes dopamine release in the NAc

Once we had established that optogenetic activation of LS-NAc, but not LS-BNST, neurons supports ICSS, we next sought to determine how activation of this projection population might facilitate positive reinforcement. The LS is a largely GABAergic structure that sends long range inhibitory projections to its downstream targets^20^. Thus, it is unclear how activation of its projections to the NAc might influence dopamine activity in the NAc to reinforce behaviors. It has been suggested that the LS, specifically the dorsal LS, increases the activity of dopaminergic neurons in the ventral tegmental area (VTA) by inhibiting tonically active GABAergic VTA neurons^41^. In order to ascertain if LS-NAc neurons might also send collaterals to the VTA, we first used a retrograde viral tracing strategy to label LS-NAc and LS-VTA neurons within the same mouse. We injected a retrogradely transported Cre virus into the NAc and a Cre-dependent GFP into the LS, followed by a retrogradely transported Flpo virus into the VTA and a Flp-dependent virus labeled with mCherry into the LS (Figure 6A). This allowed us to visualize LS-NAc neurons in green, LS-VTA neurons in red, and those that projected to both the NAc and VTA in yellow (Figure 6B). Unlike in our previous retrograde tracing experiments examining LS-NAc and LS-BNST projections where there were very few dual-labeled neurons (Figure 3C,D), we found that a subset of LS-NAc neurons also send collaterals to the VTA (Figure 6C, n = 1 mouse).

**Figure 6.**
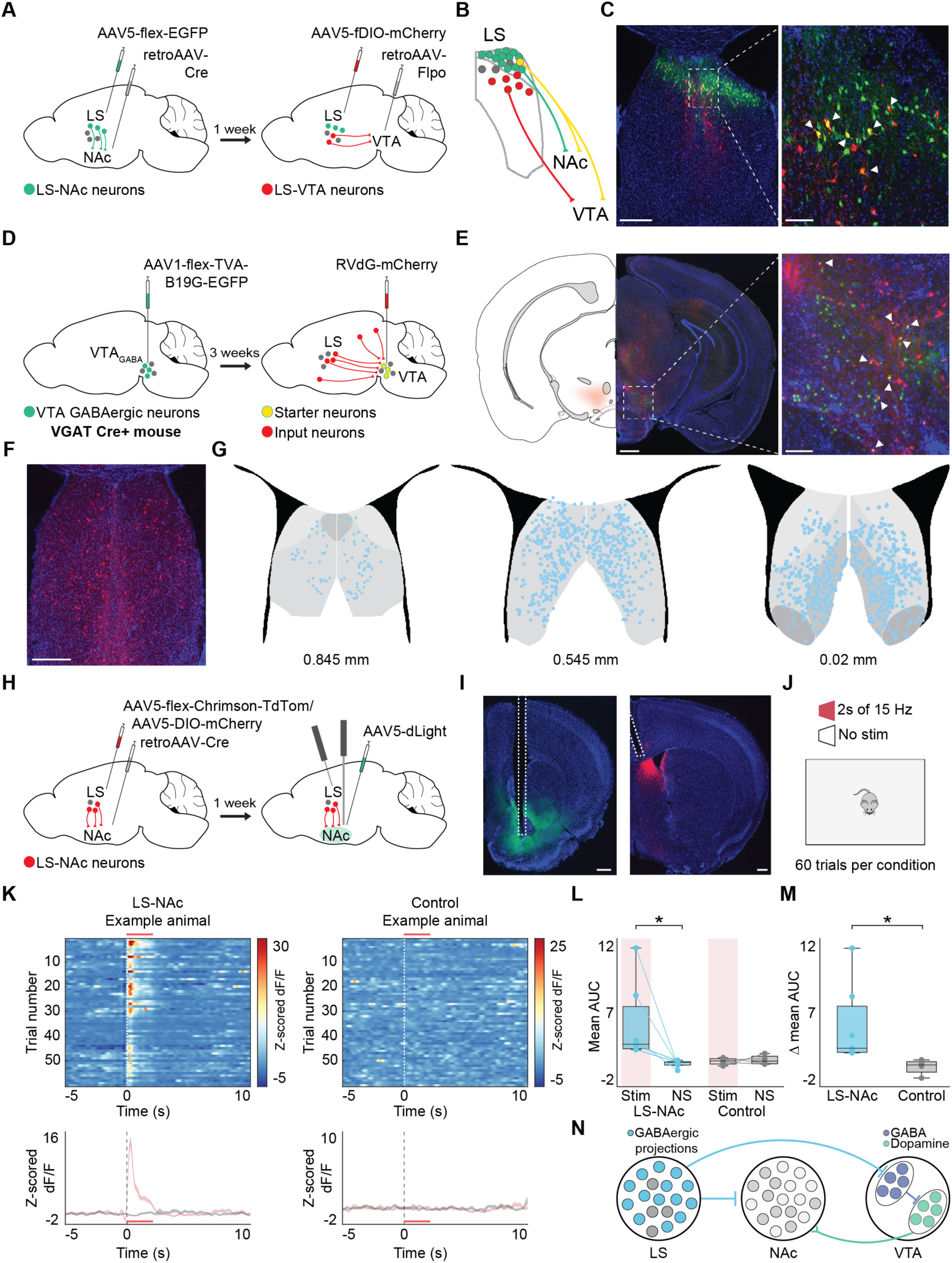
Optogenetic activation of LS-NAc neurons stimulates dopamine release in the nucleus accumbens through putative collaterals to the ventral tegmental area. A) Schematic of the viral strategy used to label LS neurons that project to the NAc and those that project to the ventral tegmental area (VTA) in the same mouse. A retrogradely transported Cre virus is injected into the NAc and a Cre-dependent GFP is injected into the LS. Then a retrogradely transported Flpo virus is injected into the VTA and a Flp-dependent mCherry in the LS. B) Depiction of viral labeling strategy whereby LS-NAc neurons are labeled with GFP (green), LS-VTA neurons are labeled with mCherry (red), and LS neurons that project to both the NAc and VTA appear yellow from dual labeling with the GFP and mCherry fluorophores. C) Representative images of LS-NAc (GFP) and LS-VTA (mCherry) neurons at 4x (left panel) and 20x (right panel) magnification. GFP expression shown in green, mCherry expression shown in red and DAPI expression shown in blue. Inset shows area outlined by dashed white square at higher magnification. Double labeled neurons appear yellow and are indicated by the filled arrowheads. Scale bars represent 250 μm and 50 μm, respectively. N = 1 mouse. D) Schematic of the modified rabies tracing strategy to label monosynaptic inputs into GABAergic VTA neurons with mCherry. A Cre-dependent helper virus labeled with GFP is injected into the VTA in a VGAT Cre+ transgenic mouse line, followed several weeks later by a modified rabies virus labeled with an mCherry fluorophore. This labels all monosynaptic inputs to the GFP-labeled VGAT VTA neurons with mCherry. E) Representative images of starter neurons in the VTA at 4x magnification. Starter neurons are neurons that have been labeled with both the helper virus (GFP) and rabies virus (mCherry). GFP expression shown in green, mCherry expression shown in red and DAPI expression shown in blue. Inset shows area outlined by dashed white square at higher magnification. VTA starter neurons labeled with both GFP and mCherry appear yellow and are indicated by the filled arrowheads. Scale bars represent 250 μm and 100 μm, respectively. F) Representative image of monosynaptic input neurons labeled with mCherry in the LS at 4x magnification. mCherry expression shown in red and DAPI expression shown in blue. Scale bar represents 250 μm. G) Coronal sections depicting LS input neurons onto GABAergic VTA neurons (individual blue dots) across 3 different A-P coordinates (0.845 mm, 0.545 mm, and 0.02 mm, all relative to bregma) mapped onto the Allen Common Coordinate framework. N = 2 mice. H) Schematic of the viral strategy used to label LS-NAc neurons with the red-shifted excitatory opsin, Chrimson (experimental, LS-NAc: n = 7 mice) or mCherry (control, n = 4 mice), and the NAc with the dopamine sensor, dLight. All mice were also implanted with a stimulating optic ferrule in the LS and a recording fiberoptic cannula in the NAc. I) Representative images of dLight labeling in the NAc (left panel) and Chrimson labeling of LS-NAc neurons in the LS (right panel) at 4x magnification. dLight expression shown in green, Chrimson expression shown in red and DAPI expression shown in blue. Tract from fiberoptic cannula (left panel) and optic ferrule (right panel) outlined in white. Scale bars represent 250 μm. J) Optogenetic activation paradigm in which a mouse in an open behavioral arena receives 2s of 15 Hz red light stimulation (stim trials) or 2s of no stimulation (NS trials) over 60 trials per stimulation condition. K) Normalized (z-scored) dLight fluorescence (dF/F) recorded in the NAc over a 15s time window during which there is 2s of red light stimulation (red bar) in a representative experimental LS-NAc mouse (left panel) and control mouse (right panel). Top panel shows a heatmap of normalized dopamine fluorescence (z-scored dF/F) on each stimulated trial aligned to trial onset and bottom panel shows average fluorescence traces ± SEM on stimulated (red) and non-stimulated (grey) trials. Dashed line at zero indicates trial onset. Red bar corresponds to 2s of red light stimulation. L) Average area under the curve (AUC) calculated in a 2.2s window following trial onset across 60 trials per stimulation condition in LS-NAc (blue) and control (grey) mice on stimulated (shaded red box) and non-stimulated trials. LS-NAc mice show an increase in average AUC on stimulated trials compared to non-stimulated trials, while control mice do not. Paired t test (LS-NAc: p = 0.0414, controls: p = 0.748). M) Change in average AUC between stimulated and non-stimulated trials in LS-NAc (blue) and control (grey) mice. LS-NAc mice show a significantly greater change in average AUC between stimulated and non-stimulated trials than controls. Welch’s t test (p = 0.0369). N = 11 mice, LS-NAc: n = 7 mice, control: n = 4 mice. *p<0.05. Boxplots: center line denotes median, box edges indicate the lower (25th) and upper (75th) quartiles, and whiskers extend to ±1.5*IQR. Individual colored points correspond to individual mice. N) Proposed circuit diagram by which optogenetic activation of LS-NAc neurons (blue) can stimulate dopamine release in the NAc through collaterals to GABAergic VTA neurons (periwinkle) and disinhibition of VTA dopamine neurons (teal).

While prior work has suggested that LS neurons may project onto GABAergic, as well as dopaminergic, neurons in the VTA, it is unclear if these monosynaptic connections to GABAergic neurons exist^41,42^. We used a modified rabies tracing strategy to confirm that LS neurons make monosynaptic connections onto GABAergic neurons in the VTA. We injected a Cre-dependent helper virus labeled with GFP into the VTA in VGAT Cre+ transgenic mice (Figure 6D). Several weeks later, we injected a modified rabies virus labeled with mCherry into the VTA of the same mice (Figure 6D). We first confirmed the presence of starter neurons (those that are labeled with both the helper virus and rabies virus) in the VTA of these mice (Figure 6E). Next, we used Wholebrain software to map the mCherry-labeled input neurons along the A-P axis (from 1.3 mm to 0.02 mm anterior to the bregma) of the LS (Figure 6F,G)^36^. We identified an average of 2,786 ± 576 total mCherry-labeled input neurons in the LS (n = 2 mice), spanning various anatomical axes, including the D-V axis (Figure 6G).

Finally, we used a combination of optogenetics and fiber photometry to test how activation of LS-NAc neurons affects dopamine release in the NAc. Using a retrogradely transported Cre virus and a Cre-dependent red-shifted excitatory opsin (Chrimson) or mCherry, we labeled LS-NAc neurons with Chrimson (experimental, n = 7 mice) or mCherry (control, n = 4 mice) and then recorded the fluorescence of a genetically encoded dopamine sensor (dLight) in the NAc (Figure 6H,I). We either stimulated LS-NAc neurons with 2s of 15 Hz red light (stimulated trials) or did not stimulate them (NS, non-stimulated trials) over 60 trials per condition as mice ambulated freely in an open behavioral arena (Figure 6J). We compared the average z-scored change in fluorescence (dF/F) of dLight in the NAc between stimulated and non-stimulated trials and between control and experimental mice (Figure 6K). We found that dopamine release, as measured by dLight fluorescence, was significantly greater in a 2.2s period following trial onset in experimental mice during stimulated trials relative to non-stimulated trials, whereas there was no significant difference in dLight fluorescence between stimulated and non-stimulated trials in the control mice (Figure 6L; paired t test, LS-NAc: p = 0.0414, control: p = 0.748). Furthermore, the change in dLight fluorescence between stimulated and non-stimulated trials (calculated as the mean AUC of dLight fluorescence on non-stimulated trials subtracted from that on stimulated trials) is significantly greater in experimental mice when compared to control mice (Figure 6M; Welch’s t test, p = 0.0369). Our findings support a circuit model in which LS-NAc activation disinhibits VTA dopamine neuron activity and increases dopamine release in the NAc via collaterals it makes onto GABAergic VTA neurons (Figure 6N).

## Discussion

In this study, we characterized the reward responses of LS neurons along its dorsoventral (D-V) axis using cellular resolution calcium imaging on a sucrose operant assay. We found that there is a gradient of reward representations such that a greater proportion of dorsal LS (LSd) neurons are excited by sucrose reward, while a greater proportion of ventral LS (LSv) neurons are inhibited by sucrose reward. Two separate projection populations within the LS occupy opposite ends of this gradient: one to the nucleus accumbens (NAc), located in the LSd, and the other to the bed nucleus of the stria terminalis (BNST), located in the LSv. Imaging of these two projection populations demonstrated that the majority of LS-NAc neurons are excited by reward and the majority of LS-BNST neurons are inhibited by reward. Additionally, activation of LS-NAc neurons and inhibition of LS-BNST neurons promotes intracranial self-stimulation (ICSS). Finally, optogenetic activation of LS-NAc neurons increases dopamine release in the NAc. These findings identify a substrate for positive reinforcement within the LS.

It has been shown that various types of information are organized along the D-V axis of the LS. For example, kin information is organized such that neurons located more dorsally in the LS respond to non-kin conspecifics, while neurons located more ventrally in the LS respond to kin^13^. Spatial and directional information is also organized along various anatomical axes of the LS^15^. Furthermore, hippocampal inputs to the LS are topographically organized and are thought to convey different streams of context-related information^43–45^. Here, we sought to localize the neural substrate of reward within the LS on a sucrose operant assay (Figure 1). We observed a gradient of reward representations such that more dorsal regions of the LS contained a larger proportion of neurons that were excited in response to sucrose reward while more ventral regions contained a larger proportion of neurons that were inhibited in response to sucrose reward (Figure 2). This dorsoventral organization of reward representations in the LS resembles that of other types of information that has been observed in the LS.

The LS is anatomically structured such that different cell types occupy various anatomical axes^20,24,25,29^. These clusters seem to correspond to the observed gradient of various types of information processing. However, specific cell types within the LS still display functional heterogeneity and their activity does not necessarily correlate with distinct behavioral outcomes^46,47^. To explain this heterogeneity, it has been proposed that there are parallel circuits of intermingled cell types within the LS that inhibit each other to funnel information to select downstream regions and promote certain behaviors^12^. It was also recently shown that subtypes of LS neurons receive preferential innervation from hippocampal and hypothalamic subregions, which convey different types of threat-related information during active avoidance behavior^25^.

In previous work, we showed that LS projection populations are anatomically segregated within the LS and that they receive different whole brain inputs, especially from the hippocampus^37^. We chose to examine the structure and function of individual LS projection populations because the anatomical organization of projection populations, as well as information encoding, along its D-V axis suggest that the LS may select downstream regions to receive separate streams of information. In particular, we focused on two projection populations, the LS-NAc and LS-BNST, which occupy the LSd and LSv, respectively. Within individual mice, we found that LS projections to the NAc and BNST are largely non-overlapping and occupy discrete LS subregions (Figure 3). Additionally, electrophysiological recordings of LS projection neurons revealed that both types produce calcium (Ca2+) spikes^39^, but that LS-NAc neurons generate them at lower current intensities than LS-BNST neurons (Figure 3). Ca2+ spikes have previously been studied in the inferior olive and cerebellum for their role in generating the plasticity that underlies motor learning, as well as oscillatory rhythms^48,49^. Consequently, the lower current threshold for Ca2+ spiking in LS-NAc neurons indicates that they require lower levels of input drive than LS-BNST neurons to enter this state of plasticity, which may differentially shape how the two populations respond to reward-related input.

Some of the earliest work mapping LS projections throughout the hypothalamus posited that each projection facilitated different motivated behaviors depending on the hypothalamic nuclei to which it projected^16^. Several studies have confirmed the importance of downstream brain regions in governing the function of the LS neurons that project to them. In one such study, neurotensin (Nts)-expressing neurons in the LS were shown to mediate homeostatic or hedonic feeding based on the hypothalamic nuclei to which they projected^50^. Additionally, somatostatin (Sst)-expressing neurons in the LSd regulate context-related avoidance behaviors and a subset of these neurons that project to the lateral hypothalamus regulate context-related feeding^47,51^. Here, we examined the reward responses of two LS populations that project to regions outside of the hypothalamus. We found that LS-NAc neurons are largely excited by sucrose reward, while LS-BNST neurons are largely inhibited by sucrose reward (Figure 4). Thus, LS projection populations that target regions outside the hypothalamus also seem to differ in function according to their downstream target, and this difference is apparent in how they encode reward.

For several decades, the site of self-stimulation within the LS has remained elusive. After finding that these two LS projection populations encoded reward differently, we examined their ability to support ICSS. We showed that optogenetic activation of LS-NAc, but not LS-BNST, neurons promotes ICSS over the course of several behavioral sessions (Figure 5). Conversely, optogenetic inhibition of LS-BNST, but not LS-NAc, neurons also elicits ICSS (Figure 5). Therefore, two opposite manipulations in two anatomically distinct populations converge on the same behavioral outcome, consistent with their opposing reward representations. We then investigated how LS-NAc neurons may mediate reward-seeking behavior and found that optogenetic activation of LS-NAc neurons stimulates dopamine release in the NAc through a putative circuit whereby LS-NAc neurons send collaterals to GABAergic VTA neurons and disinhibit dopaminergic VTA neurons (Figure 6). Given that a subset of LS-NAc neurons send collaterals to the VTA, and that LS neurons make monosynaptic contacts onto GABAergic VTA neurons, LS-NAc activation may increase dopamine release by disinhibiting VTA dopamine neurons, which places the LS-NAc population upstream of the mesolimbic dopamine system. Since the LS is a principal recipient of hippocampal output, this pathway provides a route by which contextual information can ultimately influence reward-seeking^41^.

These findings raise several compelling questions that require further exploration, especially pertaining to the dorsoventral gradient of reward representations and how it is established within the LS. In addition to the input topography discussed above, this gradient may arise from reciprocal intraseptal collaterals, such as those between the LSd and LSv that regulate aggressive behaviors^11,52^. Similar intraseptal connectivity may play a role in the different reward representations and ICSS findings observed in the LS-NAc and LS-BNST projection populations. Alternatively, this gradient may result from the inherent attributes of neurons as determined by their developmental lineage, which has been suggested for a subset of LS neurons^53,54^. Additional characterization of the cell types that constitute the LS projections to the NAc and BNST will help to clarify how these neurons function and interact with each other. Such a combined approach of cell- and projection-type specificity may be necessary moving forward to elucidate the exact role of LS neurons given the importance of the downstream region in shaping the function of LS neurons.

Along with the dorsoventral gradient of reward representations in the LS, we found that LS projection neurons display several characteristics of plasticity, namely in the generation of calcium spikes and behavioral changes with optogenetic manipulation. Similar encoding differences between the dorsal and ventral regions of the largest input to the LS, the hippocampus, have been identified on an associative learning task and these representations were found to change with learning^45^. Future work should examine if such dynamic encoding of reward is also observed in these two anatomically distinct LS projection populations. Additionally, LS neurons have been shown to modulate their activity based on motivational state and reward uncertainty^55,56^. Given that the activation of LS-NAc neurons stimulates dopamine release in the NAc, likely through collaterals to the VTA, it would be intriguing to investigate how the activity of LS projection populations shapes dopamine responses during varying states of motivation and reward availability.

This study identifies a dorsoventral gradient within the LS by which sucrose reward is encoded. It causally implicates LS projections to the NAc and BNST in reward-seeking behavior, specifically in supporting ICSS through opposing manipulations, and demonstrates that the LS-NAc population modulates dopamine release in the mesolimbic reward system. Together, these findings localize the elusive substrate of self-stimulation within the LS to discrete projection populations.

## Supplementary Figures

**Supplementary Figure 1.**
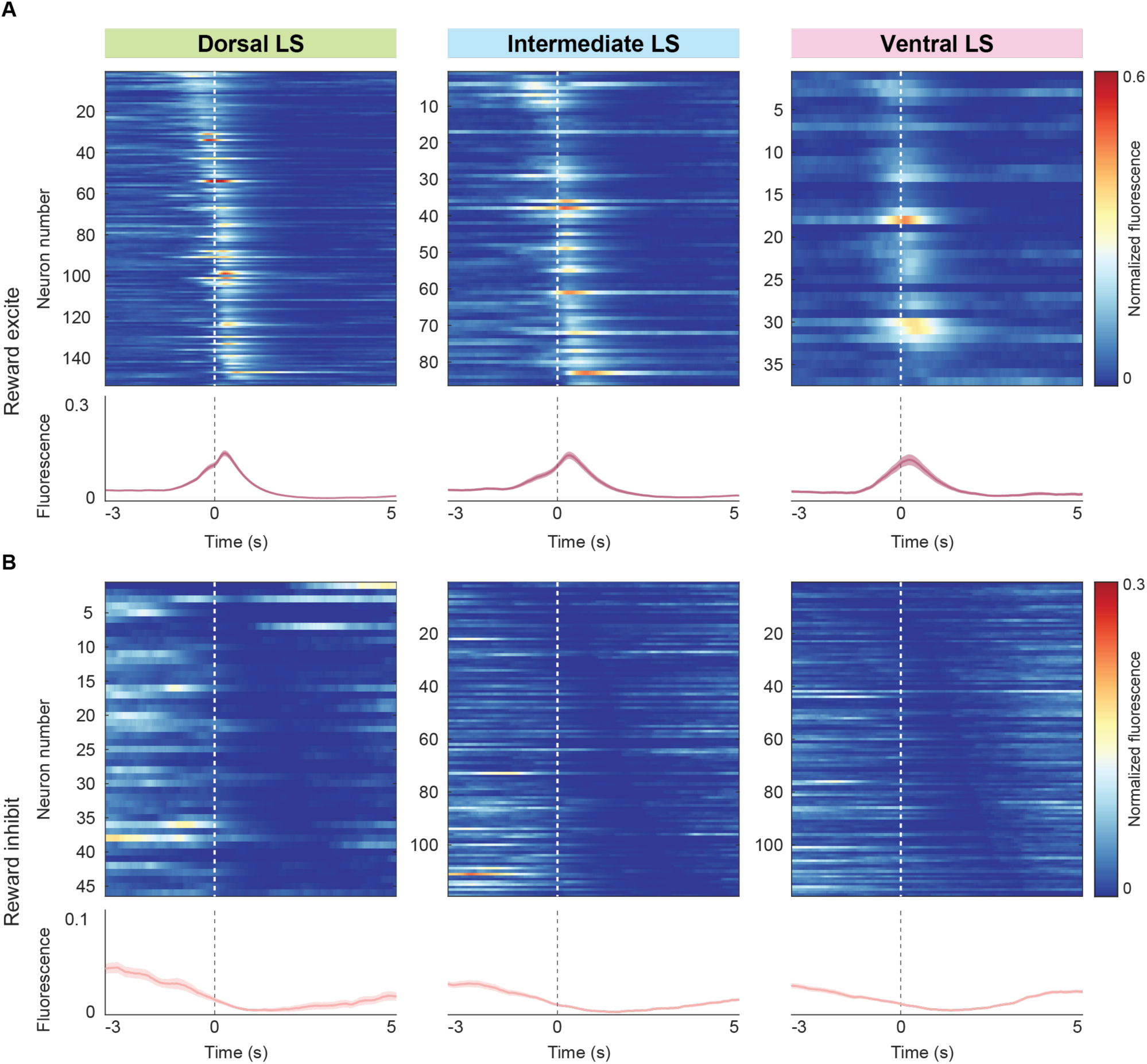
LS neurons display different proportions of reward responses along its D-V axis on the final session of a sucrose operant assay. A) Top row: Heatmaps of average normalized fluorescence of all LS neurons that are significantly positively modulated by sucrose reward consumption (reward excite) in the dorsal (left panel), intermediate (middle panel) and ventral (right panel) LS. Each row corresponds to an individual neuron. Neurons are sorted by the time of maximum fluorescence across each heatmap. Bottom row: Average normalized fluorescence traces of the neurons from the corresponding heatmap. Shaded error regions indicate ± SEM. Dashed line at zero indicates sucrose consumption onset. B) Top row: Heatmaps of average normalized fluorescence of all LS neurons that are significantly negatively modulated by sucrose reward consumption (reward inhibit) in the dorsal (left panel), intermediate (middle panel) and ventral (right panel) LS. Each row corresponds to an individual neuron. Neurons are sorted by the time of minimum fluorescence across each heatmap. Bottom row: Average normalized fluorescence traces of the neurons from the corresponding heatmap. Shaded error regions indicate ± SEM. Dashed line at zero indicates sucrose consumption onset.

**Supplementary Figure 2.**
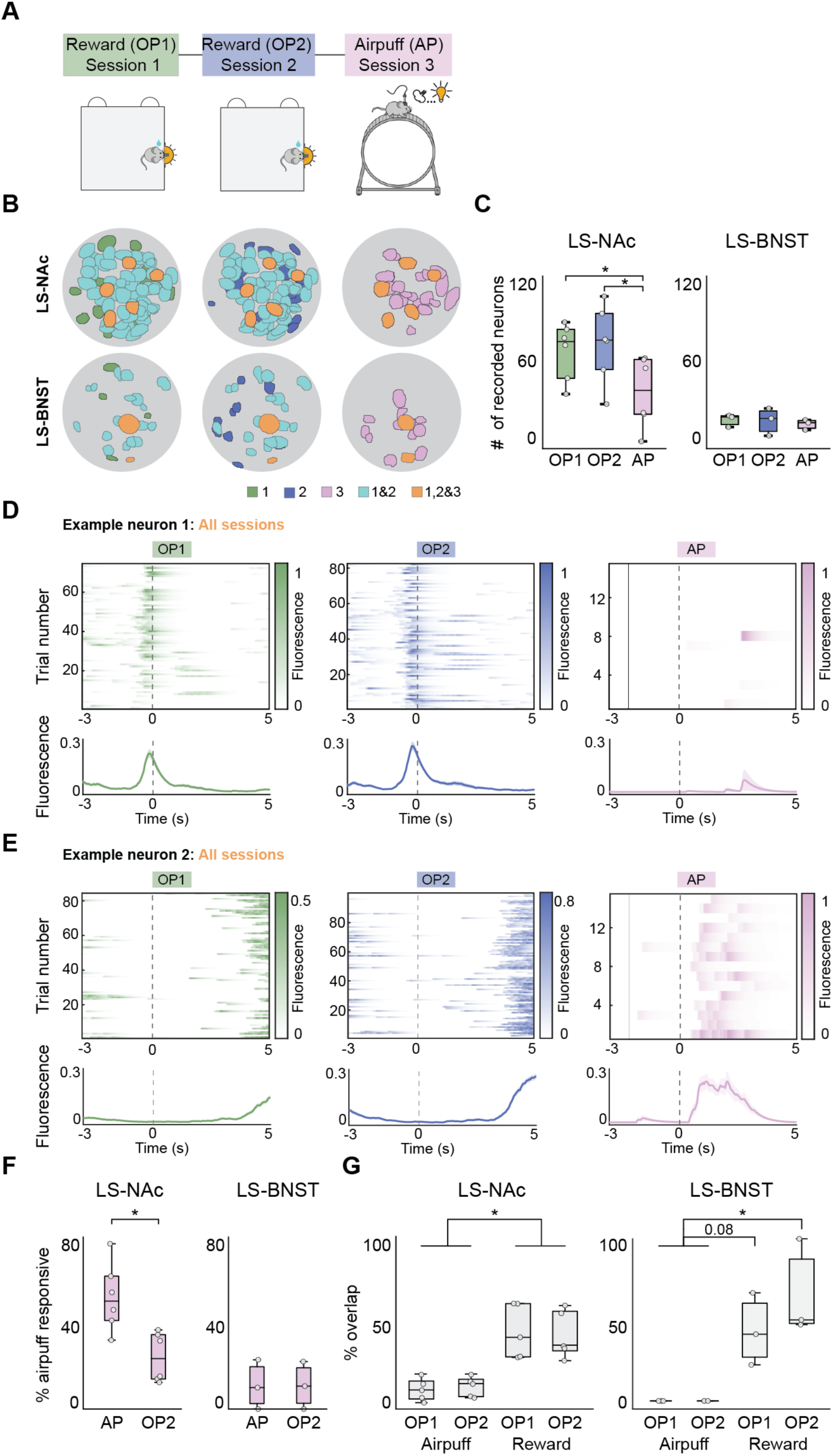
Largely non-overlapping subpopulations of LS projection neurons represent aversive and rewarding stimuli. A) Schematic of behavioral paradigms. LS-NAc and LS-BNST neurons were recorded during several behavioral paradigms, including the sucrose operant assay (OP1, left panel), an additional session of the sucrose operant assay (OP2, middle panel) and then on an aversive stimulus exposure paradigm (AP, right panel) where mice are head-fixed and receive airpuffs at a fixed interval over 20 trials. B) Imaged LS-NAc (top row) and LS-BNST (bottom row) neurons from representative mice were tracked across behavioral sessions using CellReg software^40^. The field of view from each behavioral session is shown with individual neurons colored by the sessions in which they are identified with light blue corresponding to neurons that were identified in both sucrose operant assay sessions and orange corresponding to neurons that were identified in all three behavioral paradigms. C) A greater number of LS-NAc neurons (left panel, n = 6 mice) were recorded during the sucrose operant assay sessions than the airpuff session, while a similar number of LS-BNST neurons (right panel, n = 3 mice) were recorded across all three behavioral sessions. One-way ANOVA with post-hoc t tests (LS-NAc: p = 1.60*10^-4, LS-BNST: p = 0.846). D) Normalized fluorescence of an example neuron tracked across all three behavioral sessions that is modulated by reward in the sucrose operant assay sessions (left and middle panels), but is not modulated by airpuff (right panel). Top panel shows a heatmap of normalized fluorescence on each trial aligned to stimulus onset (dashed line at zero) and bottom panel shows average fluorescence traces with shaded regions indicating ± SEM. E) Normalized fluorescence of an example neuron tracked across all three behavioral sessions that is not modulated by reward in the sucrose operant assay sessions (left and middle panels), but is modulated by airpuff (right panel). Top panel shows a heatmap of normalized fluorescence on each trial aligned to stimulus onset (dashed line at zero) and bottom panel shows average fluorescence traces with shaded regions indicating ± SEM. F) A greater proportion of LS-NAc neurons (left panel) are modulated by airpuff as a fraction of neurons recorded during the airpuff session as opposed to as a fraction of neuron recorded during second operant assay session, while a similar proportion of LS-BNST neurons (right panel) are modulated by airpuff as a fraction of recorded neurons in both airpuff and operant assay sessions. G) The proportion of airpuff modulated neurons that are also reward responsive in either the first (OP1) or second (OP2) operant assay sessions is significantly less than the reward responsive neurons that remain reward responsive across both operant assay sessions in LS-NAc (left panel) and LS-BNST mice (right panel). One-way ANOVA with post-hoc t tests (LS-NAc: p = 1.31*10^-4, LS-BNST: p = 0.00546). For post-hoc t tests see Supplementary Table 1. N = 5 LS-NAc mice (one excluded with outlier analysis), 3 LS-BNST mice. *p<0.05. Boxplots: center line denotes median, box edges indicate the lower (25th) and upper (75th) quartiles, and whiskers extend to ±1.5*IQR. Individual points correspond to individual mice.

**Supplementary Figure 3.**
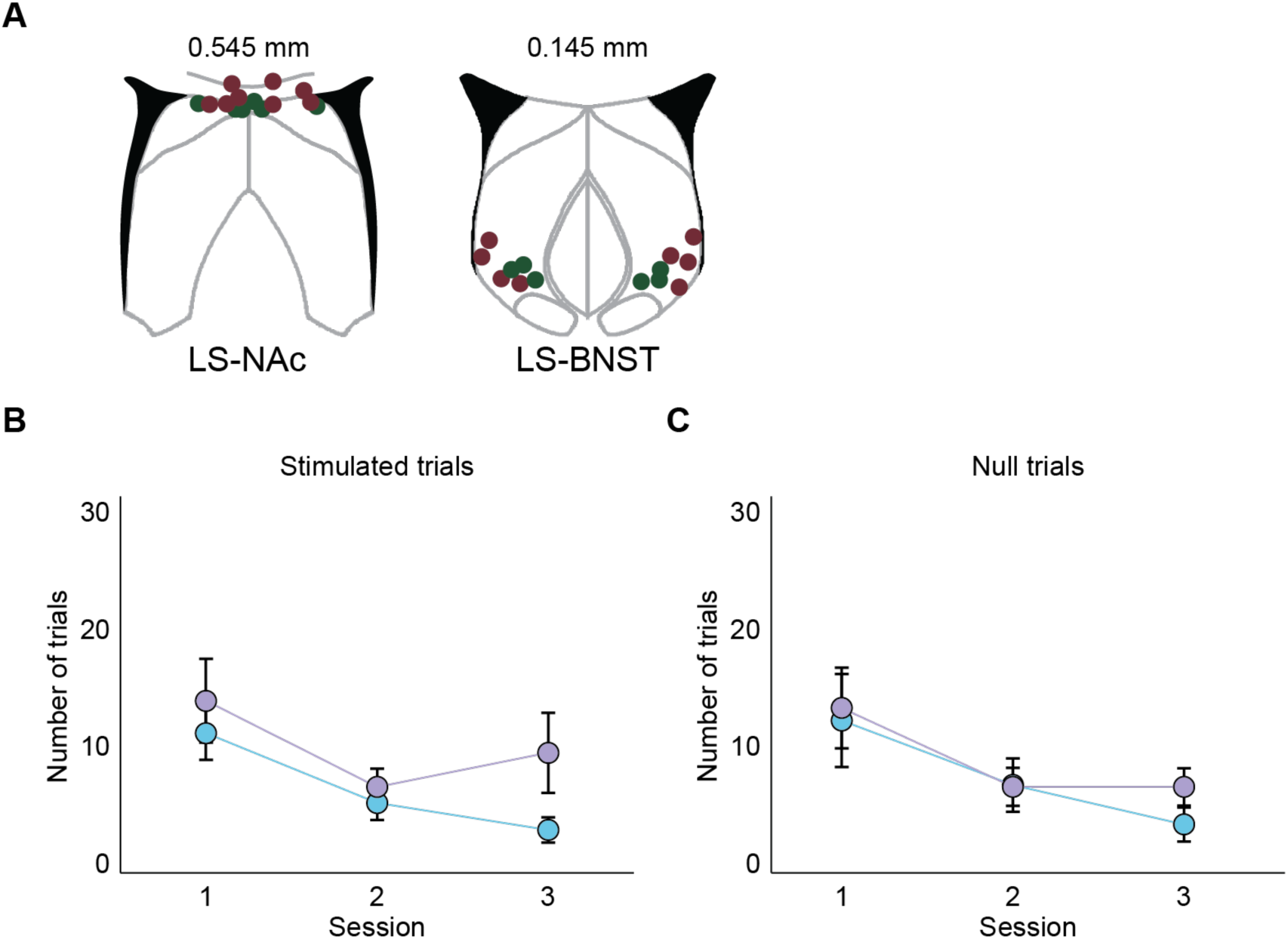
LS-NAc and LS-BNST controls do not differ in behavioral outcomes on the ICSS behavioral paradigm. A) Reconstruction of optic ferrule placement in control LS-NAc (left panel) and LS-BNST (right panel) mice. Each colored dot in the reconstruction corresponds to a ferrule in an individual mouse (dark red are mCherry mice and dark green are GFP mice) and shows the position of the ferrule relative to bregma in the Allen Mouse Brain Common Coordinate Framework. B) Control LS-NAc (blue) and LS-BNST (purple) mice complete a comparable number of stimulated trials over three ICSS behavioral sessions. Two-factor ANOVA with group type (LS-NAc controls/LS-BNST controls) and session number (1/2/3) as factors, interaction: p = 0.543, group type: p = 0.0791, session number: p = 0.0139). C) Control LS-NAc (blue) and LS-BNST (purple) mice complete a comparable number of null trials over three ICSS behavioral sessions. Two-factor ANOVA with group type (LS-NAc controls/LS-BNST controls) and session number (1/2/3) as factors, interaction: p = 0.806, group type: p = 0.519, session number: p = 0.0118). N = 7 LS-NAc control mice, 7 LS-BNST control mice. *p<0.05. Error bars indicate ± SEM.

**Supplementary Table 1.** *Compendium of statistical analyses*.

## Methods

### Experimental subjects

All viral tracing, imaging, and optogenetic experiments were performed on wild-type male C57BL/6J mice (Jax strain number: 000664) aged ∼8-14 weeks purchased from Jackson laboratories. The rabies tracing experiments in Figure 6 used the VGAT Cre+ transgenic mouse strain (Jax strain number: 028862). Mice were co-housed and given ad libitum access to food and water outside of behavioral training when mice were placed on a restricted water schedule. Mice were maintained on a reverse light dark cycle (12-hour light off-12-hour light on schedule) for all behavioral experiments. All experimental procedures were approved by and conducted in accordance with the Emory Institutional Animal Care and Use Committee.

### Viruses

The viruses used in this study, except for the modified rabies virus, were purchased from Addgene: AAV5-Syn-GCaMP6f-WPRE-SV40 (100837-AAV5), AAV5-pCAG-FLEX-EGFP-WPRE (51502-AAV5), AAVrg-hSyn-Cre-WPRE-hGH (105553-AAVrg), AAV5-EF1a-fDIO-mCherry (114471-AAV5), AAVrg-EF1a-Flpo (55637-AAVrg), AAV5-CAG-FLEX-GCaMP6f-WPRE-SV40 (100835-AAV5), AAVrg-EF1a-mCherry-IRES-Cre (55632-AAVrg), AAV5-EF1a-DIO-ChR2-EYFP-WPRE (20298-AAV5), AAV5-hSyn1-SIO-stGtACR1-FusionRed (105678-AAV5), AAV5-hSyn-DIO-mCherry (50459-AAV5), AAV1-synP-FLEX-splitTVA-EGFP-B19G (52473-AAV1), AAV5-Syn-FLEX-rc[ChrimsonR-tdTomato] (62723-AAV5), and AAV5-hSyn-dLight1.2 (111068-AAV5). The modified rabies virus, Rabies-deletedG-N2C-mCherry/EnvA, was purchased from Thomas Jefferson University (Dr. Christoph Wirblich).

### Stereotaxic surgeries

For non-specific LS imaging surgeries (Figures 1 and 2), mice aged ∼8-10 weeks (n = 15 male mice) were anesthetized with 1-2% isoflurane and placed in a stereotactic setup (Kopf). A microsyringe (Nanoject) was used to inject 0.75 µL of AAV5-Syn-GCaMP6f-WPRE-SV40 (Addgene, injected titer of 1.3x10^13 parts/ml) unilaterally into the LS at varying depths (dorsal: 0.1 mm anterior, 0.4 mm lateral and 2.8 mm in depth; intermediate: 0 mm anterior, 0.4 mm lateral and 2.8 mm in depth; ventral: 0 mm anterior, 0.45 mm lateral and 3.2 mm ventral; coordinates relative to bregma). Mice were then implanted with a 0.5 mm diameter, 6.1 mm length GRIN lens (Product ID:1050-004599, Inscopix) over the LS at varying depths (dorsal: 0.4 mm anterior, 0.35 mm lateral and 2.5 mm in depth; intermediate: 0.4 mm anterior, 0.4 mm lateral and 3.0 mm in depth; and ventral: 0.3 mm anterior, 0.5 mm lateral and 3.4 mm in depth). A metal cap was cemented over the lens to protect it. Mice were group housed following lens implant. 3-4 weeks following viral injection and lens implant, a baseplate (Product ID: 1050-004638, Inscopix) attached to the miniature microscope (nVista, Inscopix) was positioned ∼0.45 mm above the lens such that blood vessels and neurons were in focus. The baseplate was then cemented in place using dental cement (Metabond) and a 3-D printed cover was secured over the baseplate to protect the lens.

For viral tracing surgeries (Figures 3 and 6), mice aged ∼8-10 weeks (n = 10 male mice) were anesthetized with 1-2% isoflurane and placed in a stereotactic setup (Kopf). A microsyringe (Nanoject) was used to inject 0.3 µL of AAV5-pCAG-flex-EGFP-WPRE (Addgene, injected titer of 1.5x10^13 parts/ml) unilaterally into the LS (0 mm anterior, 0.4 mm lateral and 2.8 mm in depth) and 0.3 µL of AAVrg-hSyn-Cre-WPRE-hGH (Addgene, injected titer of 1.9x10^13 parts/ml) in either the NAc (1.3 mm anterior, 1.2 mm lateral and 4.7 mm in depth, n = 7 mice) or the BNST (0.15 mm anterior, 1.0 mm lateral and 4.3 mm in depth, n = 3 mice). One week after viral injection, the same mice were once again anesthetized with 1-2% isoflurane, placed in a stereotactic setup (Kopf) and a microsyringe (Nanoject) was used to inject 0.3 µL of AAV5-EF1a-fDIO-mCherry (Addgene, injected titer of 2.3x10^13 parts/ml) unilaterally into the LS (0 mm anterior, 0.4 mm lateral and 2.8 mm in depth) and 0.3 µL of AAVrg-EF1a-Flpo (Addgene, injected titer of 1.7x10^13 parts/ml) in either the NAc (1.3 mm anterior, 1.2 mm lateral and 4.7 mm in depth, n = 3 mice), the BNST (0.15 mm anterior, 1.0 mm lateral and 4.3 mm in depth, n = 6 mice), or the VTA (3.1 mm posterior, 0.35 mm lateral and 4.7 mm in depth, n = 1 mouse). The viral injections were spaced one week apart for optimal expression of fluorescent transgenes in both populations. The order of LS-NAc and LS-BNST targeting was counterbalanced across mice. Mice were allowed to recover for 3 weeks, to allow for sufficient viral expression, after which they were euthanized and brain tissue was collected for histological analysis or electrophysiological recordings.

For projection specific LS imaging surgeries (Figure 4), mice aged ∼8-10 weeks (LS-NAc = 6 male mice; LS-BNST = 6 male mice) were anesthetized with 1-2% isoflurane and placed in a stereotactic setup (Kopf). A microsyringe (Nanoject) was used to inject 0.75 µL of AAV5-CAG-flex-GCaMP6f-WPRE-SV40 (Addgene, injected titer of 2.1x10^13 parts/ml) unilaterally into the LS (0 mm anterior, 0.4 mm lateral and 2.8 mm in depth) and 0.5 µL of AAVrg-EF1a-mCherry-IRES-Cre (Addgene, injected titer of 1.8x10^13) in either the NAc (1.3 mm anterior, 1.3 mm lateral and 4.7 mm in depth) or the BNST (0.15 mm anterior, 1.0 mm lateral and 4.3 mm in depth). One week after viral injection, mice were implanted with a 0.5 mm diameter, 6.1 mm length GRIN lens (Product ID:1050-004599, Inscopix) over the specific LS projection population (LS-NAc: 0.4 mm anterior, 0.35 mm lateral and 2.5 mm in depth; LS-BNST: 0.25 mm anterior, 0.5 mm lateral and 3.5 mm in depth). A metal cap was cemented over the lens to protect it. Mice were group housed following lens implant. 3-4 weeks following viral injection and lens implant, a baseplate (Product ID: 1050-004638, Inscopix) attached to the miniature microscope (nVista, Inscopix) was positioned ∼0.45 mm above the lens such that blood vessels and neurons were in focus. The baseplate was then cemented in place using dental cement (Metabond) and a 3-D printed cover was secured over the baseplate to protect the lens.

For projection specific optogenetic activation surgeries (Figure 5), mice aged ∼8-10 weeks (n = 17 male mice) were anesthetized with 1-2% isoflurane and placed in a stereotactic setup (Kopf). A microsyringe (Nanoject) was used to inject 0.3 µL of AAV5-EF1a-DIO-ChR2-EYFP-WPRE (experimental, Addgene, injected titer of 1.8x10^13 parts/ml, n = 11 mice) or AAV5-pCAG-FLEX-EGFP-WPRE (control, Addgene, injected titer of 1.1x10^13 parts/ml, n = 6 mice) bilaterally into the LS (LS-NAc: 0.3 mm anterior, 0.3 mm lateral and 2.7 mm in depth; LS-BNST: 0 mm anterior, 0.45 mm lateral and 3.1 mm in depth) and 0.3 µL of AAVrg-EF1a-mCherry-IRES-Cre (Addgene, injected titer of 2.0x10^13 parts/ml) bilaterally into the NAc (1.3 mm anterior, 1.2 mm lateral and 4.7 mm in depth, n = 6 experimental and 3 control mice) or BNST (0.15 mm anterior, 1.0 mm lateral and 4.3 mm in depth, n = 5 experimental and 3 control mice). One week after viral injection, mice were then implanted with custom-made optic ferrules consisting of a 2.5 mm diameter metal ferrule sleeve (Product ID: MM-FER2003SS-3300, Precision Fiber Products) and a 0.3 mm diameter optic core (Product ID: FT300UMT, ThorLabs) bilaterally at a 15 degree angle into the LS above the LS-NAc projection population (0.35 mm anterior, 1.0 mm lateral and 2.45 mm in depth) or the LS-BNST projection population (0.25 mm anterior, 1.5 mm lateral and 2.9 mm in depth).

For projection specific optogenetic inhibition surgeries (Figure 5), mice aged ∼8-10 weeks (n = 19 male mice) were anesthetized with 1-2% isoflurane and placed in a stereotactic setup (Kopf). A microsyringe (Nanoject) was used to inject 0.3 µL of AAV5-hSyn1-SIO-stGtACR1-FusionRed (experimental, Addgene, injected titer of 2.1x10^13 parts/ml, n = 11 mice) or AAV5-hSyn-DIO-mCherry (control, Addgene, injected titer of 1.1x10^13 parts/ml, n = 8 mice) bilaterally into the LS (LS-NAc: 0.3 mm anterior, 0.3 mm lateral and 2.7 mm in depth; LS-BNST: 0 mm anterior, 0.45 mm lateral and 3.1 mm in depth) and 0.3 µL of AAVrg-hSyn-Cre-WPRE-hGH (Addgene, injected titer of 1.9x10^13 parts/ml) bilaterally into the NAc (1.3 mm anterior, 1.2 mm lateral and 4.7 mm in depth, n = 6 experimental and 4 control mice) or BNST (0.15 mm anterior, 1.0 mm lateral and 4.3 mm in depth, n = 5 experimental and 4 control mice). One week after viral injection, mice were then implanted with custom-made optic ferrules consisting of a 2.5 mm diameter metal ferrule sleeve (Product ID: MM-FER2003SS-3300, Precision Fiber Products) and a 0.3 mm diameter optic core (Product ID: FT300UMT, ThorLabs) bilaterally at a 15 degree angle into the LS above the LS-NAc projection population (0.35 mm anterior, 1.0 mm lateral and 2.45 mm in depth) or the LS-BNST projection population (0.25 mm anterior, 1.5 mm lateral and 2.9 mm in depth)

For monosynaptic rabies tracing surgeries (Figure 6), VGAT Cre+ transgenic mice aged ∼8-10 weeks (n = 2 male mice) were anesthetized with 1-2% isoflurane and placed in a stereotactic setup (Kopf). A microsyringe (Nanoject) was used to inject 0.5 µL of Cre-dependent helper virus, AAV1-synP-FLEX-splitTVA-EGFP-B19G (Addgene, injected titer of 2.4x10^13 parts/ml), unilaterally into the VTA (3.1 mm posterior, 0.35 mm lateral and 4.7 mm in depth). Three weeks after viral injection, the same mice were once again anesthetized with 1-2% isoflurane, placed in a stereotactic setup (Kopf) and a microsyringe (Nanoject) was used to inject 0.5 µL of modified rabies virus, Rabies-deletedG-N2C-mCherry/EnvA (Thomas Jefferson University, injected titer of 8x10^8 parts/ml) unilaterally into the VTA (3.1 mm posterior, 0.35 mm lateral and 4.7 mm in depth). Mice were allowed to recover for 7 days, after which they were euthanized and brain tissue was collected for histological analysis.

For combined optogenetic/fiber photometry surgeries (Figure 6), mice aged ∼8-10 weeks (n = 11 male mice) were anesthetized with 1-2% isoflurane and placed in a stereotactic setup (Kopf). A microsyringe (Nanoject) was used to inject 0.4 µL of AAV5-Syn-FLEX-rc[ChrimsonR-tdTomato] (experimental, Addgene, injected titer of 2.2x10^13 parts/ml, n = 7 mice) or AAV5-hSyn-DIO-mCherry (control, Addgene, injected titer of 1.1x10^13 parts/ml, n = 4 mice) unilaterally into the LS (0.3 mm anterior, 0.3 mm lateral and 2.7 mm in depth) and 0.4 µL of AAVrg-hSyn-Cre-WPRE-hGH (Addgene, injected titer of 1.9x10^13 parts/ml) unilaterally into the NAc (1.3 mm anterior, 1.2 mm lateral and 4.7 mm in depth, n = 7 experimental and 4 control mice). One week after the first viral injection, the same mice were injected with 0.6 µL of AAV5-hSyn-dLight1.2 (Addgene, injected titer of 2.4x10^13 parts/ml) in the ipsilateral NAc (1.3 mm anterior, 1.15 mm lateral and 4.7 mm in depth) and then implanted with a custom-made optic ferrule consisting of a 2.5 mm diameter metal ferrule sleeve (Product ID: MM-FER2003SS-3300, Precision Fiber Products) and a 0.3 mm diameter optic core (Product ID: FT300UMT, ThorLabs) unilaterally at a 15 degree angle into the LS above the LS-NAc projection population (0.5 mm anterior, 0.4 mm lateral (contralateral to the injection) and 2.3 mm in depth). At the same time, mice were implanted with a 6 mm long fiberoptic cannula with a 0.48-NA, 0.4 mm diameter optic core and a 2.5 mm black ceramic ferrule (Product ID: FOC-BF-400-250, Neurophotometrics) in the ipsilateral NAc (1.3 mm anterior, 1.15 mm lateral and 4.7 mm in depth).

### Behavioral Experiments

Mice were co-housed and placed on a restricted water schedule prior to the start of training. At the end of each day mice were provided with 1.5 mL of water in addition to their consumption during training. Mice were then sequentially trained to engage in reward-seeking behavior on an operant assay as has been previously described^57,58^.

#### Training stage 1: Operant conditioning

In the first stage (operant conditioning), mice were trained to associate poking the reward port with a sucrose reward. Nose port entry triggered the port LED light on and resulted in the delivery of a sucrose reward of 10 µL of 10% sucrose in water. Each session lasted an hour and animals were run through the assay once a day. Mice had to reach a minimum of 50 pokes in a single session to progress to the next training stage. If the animal did not engage with the reward port for more than two minutes after the start of the trial, then the reward port delivered 30 µL liquid without engaging the LED. This automatic reward delivery also occurred at the beginning of the first trial in each session to prime the animal to engage with the port.

#### Training stage 2: Single port

In the second stage (single port), mice were conditionally rewarded with 10 µL of sucrose for poking the reward port within 8s of the reward port LED turning on. Trials were separated by a 30s ITI, and the animal had to refrain from entering the port for 3s for the next trial to start. A trial was considered a sucrose reward fail if the mouse failed to enter the sucrose reward port after it had been illuminated for 8s. Mice had to reach a minimum of 40 successful trials during one session to proceed to the next training stage.

#### Sucrose operant assay

In the third stage (sucrose operant assay), the choice port LED furthest from the sucrose reward port (sucrose choice port) turned on to indicate trial start. This assay is self-paced, so mice had an indefinite amount of time to nose poke the illuminated sucrose choice port. A successful choice port poke resulted in the LED flashing on and off successively 3 times in 0.1s intervals, after which the reward port LED turned on for up to 8s. Mice had to nose poke the reward port within this window to receive a 10 µL sucrose reward. Failure to do so resulted in a reward fail and initiation of a new trial. Trials were separated by a 15s ITI. In addition, the current trial ended if the animal engaged with the other inactive port instead of the sucrose choice port during the choice period. Mice had to refrain from entering the sucrose choice port for 3s for the next trial to start. Mice were considered to be fully trained on the behavioral assay when they completed more than 70 successful sucrose trials in a session.

#### Aversive stimulus exposure paradigm

Mice were returned to ad libitum water access for several days following the sucrose operant assay imaging experiments (Supplementary Figure 2). On the day of the aversive stimulus exposure paradigm, mice were head-fixed above a custom made running wheel^59^ and the imaging scope was secured to the baseplate of the mouse. Mice were allowed to habituate to the scope and head fixation for 10 minutes prior to the start of the session. They were then subjected to 20 trials of manually delivered 2s airpuffs separated by a 20s ITI. Airpuff delivery was timed to the 2s illumination of a port LED that was positioned in front of the mouse to ensure visibility throughout the duration of the imaging session.

### Behavioral Analysis

All behavioral session videos were recorded at 40 Hz using Pylon software. The BPOD (Sanworks) data acquisition system also created a MATLAB output file containing timestamps of all behavioral metrics from each behavioral session. Following behavioral sessions, videos were analyzed and behavior was quantified. Several metrics were used to quantify mouse behavior for a given behavioral session, including number of sucrose trials, choice latency and reward latency. The number of sucrose trials was determined as the total number of times that an animal nose-poked the sucrose choice port and entered the sucrose reward port when the reward was available for consumption (8s following choice port nose-poke).

### Endoscopic Calcium Imaging Experiments and Analysis

∼3-4 week post lens implant surgery, GRIN lens (0.5 mm diameter, 6.1 mm length) implanted mice (n = 15 non-specific LS imaging mice and 12 projection specific imaging mice) were trained sequentially on the sucrose operant assay while calcium imaging recordings were performed during each session. On each day of imaging, mice were allowed to habituate to the scope for 5 minutes prior to the start of the session. During the behavioral imaging session, the LED power was set to 0.6-0.7 and the analog gain on the image sensor was set to 1.6-1.8. Images were acquired at 20 Hz using Inscopix nVista software. Following imaging acquisition, data were spatially downsampled by a factor of 4 and motion corrected using Inscopix software. The CNMFe algorithm was then used to identify individual neurons and their fluorescence traces^35^. The fluorescence traces were subsequently aligned to the behavioral data and used for task modulation analysis. Custom MATLAB software was used to analyze all the imaging data.

### Acute slice preparation

Mice were anesthetized and perfused with ice-cold cutting solution (in mM) 87 NaCl, 25 NaHCO3, 2.5 KCl, 1.25 NaH2PO4, 7 MgCl2, 0.5 CaCl2, 10 glucose, and 7 sucrose. Mice (n = 3) were then euthanized by decapitation and the brain was immediately removed by dissection. Brain slices (250 μm) were sectioned in the coronal plane using a vibrating blade microtome (VT1200S, Leica Biosystems) in the same solution. Slices were transferred to an incubation chamber and maintained at 34°C for ∼30 min and then at 23–24°C thereafter. During whole-cell recordings, slices were continuously perfused with (in mM) 128 NaCl, 26.2 NaHCO3, 2.5 KCl, 1 NaH2PO4, 1.5 CaCl2, 1.5MgCl2, and 11 glucose, maintained at 30.0°C ± 0.5°C. All solutions were equilibrated and maintained with carbogen gas (95% O2/5% CO2) throughout.

### Electrophysiology

Labeled neurons were targeted for somatic whole-cell recording in the LS by combining gradient-contrast video (Retiga ELECTRO; Teledyne) microscopy with epifluorescent illumination (XCite white light source) using a commercial (Olympus) upright microscope with a manual turret to switch between GFP and mCherry excitation/emission. Recordings were obtained using Multiclamp 700B amplifiers (Molecular Devices). Signals were filtered at 6-10 kHz and sampled at 50 kHz with a Digidata 1440B digitizer (Molecular Devices). Borosilicate patch pipettes were filled with an intracellular solution containing (in mM) 124 potassium gluconate, 2 KCl, 9 HEPES, 4 MgCl2, 4 NaATP, 3 l-ascorbic acid, and 0.5 NaGTP. Pipette capacitance was neutralized in all recordings and electrode series resistance compensated using bridge balance in current clamp. Liquid junction potentials were uncorrected. Recordings had a series resistance <20 MΩ. Membrane potentials were maintained with constant bias current near –75 mV for both neuron types (-75.6+/-1.9 vs -72.9+/-1.6 mV; p=0.27; unpaired t test; LS-BNST vs LS-NAc populations, respectively). AP trains were initiated by somatic current injection at varying levels (300 ms each). Both GFP- and mCherry-labeled cells were recorded in each slice.

### Optogenetic Activation Experiments

∼2 weeks post ferrule implants, ChR2- and GFP-expressing mice (ChR2: n = 11 mice, GFP: n = 6 mice) were run on three consecutive sessions of the optogenetic intracranial self-stimulation (ICSS) paradigm. Before each behavioral session, optic ferrules on the animal’s head were coupled to patch cords (Product ID: SBP, Doric) attached to a commutator (Product ID: FRJ_1x1_FC, Doric) using ceramic split mating sleeves (Product ID: ADAF, ThorLabs). The commutator was connected to a 473 nm laser (Product ID: BL473T8-200FC, SLOC Lasers) via a patch cable (Product ID: M72L05, ThorLabs). During each one-hour behavioral session, mice were allowed to freely choose between a port designated as the stimulation port which was paired with laser stimulation and a null port (no stimulation). Nose-poking the stimulation port resulted in 2s of blue light stimulation (473 nm, 15 Hz, 20 ms pulse duration, 10 mW at fiber tip). Each trial type (stimulation/null) was followed by a 15s ITI before the initiation of the next trial, signaled by illumination of both ports. The stimulation and null ports were counterbalanced across mice. BPOD Sanworks and a custom MATLAB program were used to control laser pulses. Behavioral analysis was performed using MATLAB software to compare between (null versus stimulated trials) and across (GFP control versus ChR2 experimental) mice.

### Optogenetic Inhibition Experiments

∼2 weeks post ferrule implants, GtACR- and mCherry-expressing mice (GtACR: n = 11 mice, mCherry: n = 8 mice) were run on one session of the optogenetic ICSS paradigm. Before each behavioral session, optic ferrules on the animal’s head were coupled to patch cords (Product ID: SBP, Doric) attached to a commutator (Product ID: FRJ_1x1_FC, Doric) using ceramic split mating sleeves (Product ID: ADAF, ThorLabs). The commutator was connected to a 532 nm laser (Product ID: GL532T3-200FC, SLOC Lasers) via a patch cable (Product ID: M72L05, ThorLabs). As with the optogenetic activation experiments, During each one-hour behavioral session, mice were allowed to freely choose between a port designated as the stimulation port which was paired with laser stimulation and a null port (no stimulation). Nose-poking the stimulation port resulted in 2s of green light stimulation (532 nm, 2s continuous stimulation, 12 mW at fiber tip). Each trial type (stimulation/null) was followed by a 15s ITI before the initiation of the next trial, signaled by illumination of both ports. The stimulation and null ports were counterbalanced across mice. BPOD Sanworks and a custom MATLAB program were used to control laser pulses. Behavioral analysis was performed using MATLAB software to compare between (null versus stimulated trials) and across (mCherry control versus GtACR experimental) mice.

### Combined Optogenetic/Fiber Photometry Experiments and Analysis

∼3 weeks post optic ferrule and fiberoptic cannula implants, Chrimson- and mCherry-expressing mice (Chrimson: n = 7 mice, mCherry: n = 4 mice) were run on three consecutive sessions of red-light optogenetic stimulation and simultaneous dopamine fluorescence fiber photometry recordings. Before each behavioral session, the optic ferrule on the animal’s head was coupled to a patch cord (Product ID: MFP, Doric) that was attached to a commutator (Product ID: FRJ_1x1_FC, Doric) using a ceramic split mating sleeve (Product ID: ADAF, ThorLabs). The commutator was connected to a 635 nm laser (Product ID: RLM635TA-200FC, SLOC Lasers) via a patch cable (Product ID: M72L05, ThorLabs). The fiberoptic cannula was also coupled to a low autofluorescence patch cord (Product ID: MFP-LAF, Doric) using a ceramic split mating sleeve (Product ID: ADAF, ThorLabs). The patch cord was directly connected to a Doric four port fluorescence mini cube, which consisted of a 465 nm excitation channel (fluorescence channel) and a 405 nm isosbestic channel (control channel). During each behavioral session, mice were either stimulated (635 nm, 15 Hz, 20 ms pulse duration, 10 mW at fiber tip) or not stimulated for 2s followed by a 20s ITI over the course of 40 trials (20 stimulated, 20 non-stimulated) per session for three sessions. Fluorescence signals were acquired at 60 Hz using a Doric fiber photometry console. The BPOD Sanworks and a custom MATLAB program were used to control laser and acquisition pulses. Following fiber photometry recordings, data were imported to Doric Neuroscience Studio for analysis and smoothed using a low-pass Butterworth filter. The isosbestic channel was then regressed out from the fluorescence channel to control for motion artifacts and photobleaching. The fitted fluorescence traces (dF/F) were then exported, aligned to BPOD behavioral data in MATLAB and z-scored using a previously described method in which baseline fluorescence traces were subtracted from fluorescence traces aligned to the behavioral event and divided by the baseline standard deviation^60^. The baseline fluorescence trace for each behavioral event was defined as that from a 5s window in the middle (7.5s to 12.5s) of the 20s ITI following the behavioral event. The aligned z-scored fluorescence traces were averaged across 60 trials per stimulation condition. Area under the curve (AUC) analysis was performed on z-scored fluorescence from each trial in a 2.2s window following trial start using the *trapz* function in MATLAB.

### Histology and Microscopy

After the completion of imaging, viral tracing, and optogenetic experiments, mice were sacrificed and perfused with 0.5% PBS followed by 4% paraformaldehyde. The brains were extracted and fixed in 4% PFA overnight at 4°C. They were then transferred to 30% sucrose solution and allowed to fix for a minimum of 24 hours. To visualize viral expression and GRIN lens/optic ferrule/fiberoptic cannula placement, brains were sectioned into 50 µm slices using an Epredia HM430 microtome and the slices containing the target brain regions were mounted in DAPI solution (Product ID: 0100-20, Southern Biotech). Slices were imaged using fluorescence microscopy (Keyence BZ-X800). Images from viral tracing were registered using ABBA and exported to QuPath for cell counting and quantification^38,61^. All other images were registered using WholeBrain software^36^ to determine GRIN lens and optic ferrule placement.

### Data Analysis and Statistics

#### Task modulation analysis

We used a previously described method to classify neurons as significantly modulated by various task events (sucrose choice and reward)^57^. To do this, we examined the normalized change in fluorescence in each peri-behavioral task window. Given the temporal proximity of task events (especially choice and reward delivery), we designated a 1s window around each task event and calculated the mean fluorescence within that window. Peri-behavioral task windows were assigned as follows to eliminate overlap between task event responses: trial start: 0.5s before and after; choice: 0.05s before and 0.95s after; reward: 0.05s before and 0.95s after. For excitatory responses, we compared the maximum value of the mean fluorescence to the maximum value expected by a shuffled distribution (>99.5 percentile). For inhibitory responses, which require a sustained suppression of fluorescence, we determined if the mean value of fluorescence across the time window was lower than that expected in a shuffled distribution (<0.5 percentile). The shuffled distribution was generated by circularly shifting (coded in MATLAB using *circshift*) individual fluorescence traces by random values over 1000 iterations and then averaging shuffled activity within each peri-behavioral time window.

#### Whole brain registration

To determine GRIN lens (Figures 1, 2 and 4) and optic ferrule placement (Figure 5), as well as for the identification of LS neurons that project to the VTA in the rabies tracing experiments (Figure 6D-G), we used an open-source Wholebrain R package^36^. Images of coronal sections were cropped and converted into grayscale 8-bit TIFF files in order to be compatible with the Wholebrain package. Each section was manually assigned to an A-P coordinate and registered to the corresponding Allen Brain Atlas section using Wholebrain software whereby individual points are referenced between the tissue section and the Allen Brain Atlas section to transform it accordingly^36^. For GRIN lens and optic ferrule placement, reference points were added to denote the location of the lens or ferrule. For monosynaptic rabies tracing, a segmentation filter was applied to identify fluorescently-labeled neurons within the LS based on brightness and soma size. Individual neurons were automatically detected and manually confirmed in each section.

#### ABBA registration and cell counting

For viral tracing experiments (Figure 3), we used ABBA software to register individual coronal brain sections to the Allen Brain Atlas^38^. Images of 50 µm coronal sections at 3 manually assigned A-P coordinates (0.845 mm, 0.545 mm and 0.02 mm) were uploaded as TIFF files to ABBA through the Fiji plugin. Each section was then aligned to an Allen Brain Atlas section at each A-P coordinate and the ‘Elastix’ tool was used to perform affine registration of the tissue section to the corresponding Allen Brain Atlas section. The ‘Big Warp’ tool was used to further refine the registration based on anatomical landmarks such as the corpus callosum and lateral ventricles. Registered sections were then exported to QuPath, where fluorescently labeled LS neurons were automatically detected and manually confirmed using its ‘Cell Detection’ analysis tool (in both red and green channels) within the annotated anatomical boundaries determined through the registration process^61^.

#### Density contour plots

We created density contour plots of LS-BNST and LS-NAc neurons at 3 different A-P coordinates (0.845 mm, 0.545 mm and 0.02 mm) that span the entirety of the LS (Figure 3E) by concatenating projection neurons across animals (n = 6 mice) and calculating the density values from 0.3 to 0.9 (coded in MATLAB using *ksdensity*). Normalized contours (based on the peak density value) were then plotted and overlaid on an outline of the LS corresponding to the Allen Brain Atlas at each A-P coordinate. The centroid was depicted as the center of the 90% contour (black cross in Figure 3E and colored dot in Figure 3F). Kolmogorov-Smirnov (KS) tests were performed as pairwise comparisons between the 90% contours of the two LS projections at each A-P coordinate.

#### LS neuron distributions

To examine the distribution of the LS projection (LS-NAc and LS-BNST) neurons along the D-V and M-L axes (Figure 3G), we binned the data across its ranges (D-V axis: 0.2 to -2.0 mm and M-L axis: 0 to 1.0 mm) into approximately 240 and 110 µm bins, respectively. We then interpolated the means and standard errors of the data from each projection with the Catmull-Rom spline function utilized by Microsoft Excel.

#### Electrophysiology and classification of putative Ca2+ spikes

Na^+^ spike parameters (i.e., 1st spike) and passive properties were analysed using standard methods^62^. Putative Ca^2+^ spikes were automatically detected using a custom python pipeline built on top of pyabf (v2.3) and scipy.signal. The pipeline detected every spike-like event in each sweep, and classified each event as Na⁺, putative Ca²⁺, or undefined using a hierarchical rule. All analyses were performed offline. An event was classified as Na⁺ if its half-width was 2.0 ms or less, or if its peak voltage was 10 mV or more and its half-width was less than 3.0 ms. Putative Ca^2+^ spikes are complex events which are considered obvious by eye to experienced physiologists, but not easily designated by a single variable^39^. Thus, a putative Ca^2+^ event was classified conservatively, only if these criteria were met: a half-width of at least 5.0 ms together with a peak voltage below 5 mV; an upstroke-onset voltage above −47 mV together with a peak voltage below 10 mV and a half-width of at least 3.5 ms; or a peak voltage below −10 mV together with a maximum rising dV/dt below 15 V/s. Spikes that fell between these variables were considered undefined by the program. After automated classification, designated Ca^2+^ spikes were hand-verified by an experienced physiologist in a manner blinded to cell type.

#### Decoding projection population using neural data

To determine if the identity of the projection population (Figure 4G) could be decoded from neural activity during reward periods, we used logistic regression (coded in MATLAB using *fitclinear*) for the neural data from the corresponding reward time window (0.5s before to 5.5s after). For each mouse, we used a matched number of trials and neurons to construct a three-dimensional array of [*number of neurons x number of trials x time window of z-scored fluorescence data*]. This ensured that each binary decoder was trained with balanced labels accounting for the variability in the number of sucrose trials and neurons recorded from each mouse. The three-dimensional array was averaged across neurons to use as an input to the regression. For every sucrose trial, we trained binary decoders to predict if the averaged neural response corresponded to either the LS-NAc or LS-BNST projection population. The input to the decoder was divided into train and test sets with a 70-30 train-test split and the train set was used to optimize the regression model hyperparameters using Lasso regularization. We evaluated the performance of the decoder, (reported as decoding accuracy) by applying the trained model to predict labels for both a test set and a generalization set. Across the 6 LS-NAc and 6 LS-BNST mice, we performed pairwise comparisons and calculated decoding accuracy for 500 iterations in which we randomly sampled trials into the train and test set. The average test decoding accuracy for each animal was reported as an average of 500 iterations. To ensure that the reported decoding accuracy reflected differences in neural responses between projection populations and not individual animals, we created a generalization set in which the trained model from one pairwise comparison was applied to all other pairwise comparisons to predict projection population labels for the data that were not part of the train/test set (average decoding accuracy, Figure 4G). Finally, we calculated the decoding accuracy for a shuffled dataset that was created by shuffling the z-scored fluorescence. Average decoding accuracy is reported across all projection specific LS imaging mice.

